# Structural basis of transcription inhibition by the DNA mimic protein Ocr of bacteriophage T7

**DOI:** 10.1101/822460

**Authors:** Fuzhou Ye, Ioly Kotta-Loizou, Milija Jovanovic, Xiaojiao Liu, David T. F. Dryden, Martin Buck, Xiaodong Zhang

## Abstract

Bacteriophage T7 infects *Escherichia coli* and evades the host defence system. The Ocr protein of T7 was shown to exist as a dimer mimicking DNA and to bind to host restriction enzymes, thus preventing the degradation of the viral genome by the host. Here we report that Ocr can also inhibit host transcription by directly binding to bacterial RNA polymerase (RNAP) and competing with the recruitment of RNAP by sigma factors. Using cryo electron microscopy, we determined the structures of Ocr bound to RNAP. The structures show that an Ocr dimer binds to RNAP in the cleft, where key regions of sigma bind and where DNA resides during transcription synthesis, thus providing a structural basis for the transcription inhibition. Our results reveal the versatility of Ocr in interfering with host systems and suggest possible strategies that could be exploited in adopting DNA mimicry as a basis for forming novel antibiotics.

**Impact statement:** DNA mimicry Ocr protein, a well-studied T7 phage protein that inhibits host restriction enzymes, can also inhibit host transcription through competing with sigma factors in binding to RNA polymerase.

## Introduction

Bacteriophage T7 infects *Escherichia coli* and hijacks the host cellular machinery to replicate its genome (Studier 1972, Kruger and Schroeder 1981). The T7 genome encodes 56 proteins with many functioning as structural proteins for the bacteriophage. A number of T7 proteins are known to specifically inhibit the bacterial cellular machinery. For example, proteins gp0.7, gp2 and gp5.7 inhibit cellular transcription (Camara, Liu et al. 2010, Tabib-Salazar, Liu et al. 2018) whereas gp0.3 inhibits restriction enzymes (Studier 1975).

Gp0.3 is the first T7 gene expressed after infection and T7 variants lacking gene 0.3 were shown to have genomes susceptible to *E. coli* restriction systems (Studier 1975). Subsequently the 117 amino acid protein gp0.3 was named Overcome Classical Restriction (Ocr) (Kruger and Schroeder 1981). Ocr is abundantly expressed and forms a dimer that mimics the structure of a slightly bent 20 base pair B-form DNA (Walkinshaw, Taylor et al. 2002) (Issinger and Hausmann 1972) and blocks the DNA binding grooves of the type I restriction/modification enzyme, preventing the degradation and modification of the T7 genome by the host. Intriguingly, Type I restriction/modification enzyme is present in very low numbers (estimated at ∼60 molecules per cell, (Kelleher and Raleigh 1994)). Since Ocr is a DNA mimicry protein, it is possible that the abundantly expressed Ocr also interferes with other DNA processing systems of the host. Indeed early evidence of an interaction between Ocr and the host RNA polymerase (RNAP) was obtained using pull-down affinity chromatography (Ratner 1974).

RNA polymerase is the central enzyme for transcription, which is a highly controlled process and can be regulated at numerous distinct functional stages (Kornberg 1998, Decker and Hinton 2013). The large majority of transcription regulation, however, is executed at the recruitment and initiation stage (Browning and Busby 2004, Hahn and Young 2011, Browning and Busby 2016). To ensure transcription specificity, bacterial RNAP relies on sigma (σ) factors to recognise gene-specific promoter regions. *E. coli* has seven sigma factors which can be grouped into two classes, the σ^70^ class represented by σ^70^, responsible for transcribing housekeeping genes, and the σ^54^ class, responsible for transcribing stress-induced genes (Feklistov, Sharon et al. 2014, Browning and Busby 2016). Much work has yielded a detailed mechanistic understanding of how transcription directed by σ^70^ and σ^54^ is initiated (Zhang, Feng et al. 2012, Glyde, Ye et al. 2018). Specifically, the two large RNAP subunits β and β’ form a crab claw structure that encloses the DNA binding cleft, accommodating the transcription bubble and the downstream double-stranded (ds) DNA (Bae, Feklistov et al. 2015, Zuo and Steitz 2015).

Inhibiting host transcription is widely exploited by bacteriophages including T7 on *E. coli* and P23-45 on *T. thermophilus* (Tagami, Sekine et al. 2014, Tabib-Salazar, Liu et al. 2017, Ooi, Murayama et al. 2018). Gp0.7, gp2 and gp5.7 of T7 were shown to inhibit bacterial RNAP (Camara, Liu et al. 2010, Tabib-Salazar, Liu et al. 2017). Gp0.7 is a protein kinase that phosphorylates the *E. coli* RNAP-σ^70^, resulting in transcription termination at sites located between the early and middle genes on the T7 genome, gp2 specifically inhibits σ^70^-dependent transcription and gp5.7 inhibits σ^S^, responsible for stationary phase adaptation (Bae, Davis et al. 2013, Tabib-Salazar, Liu et al. 2018). Two proteins gp39 and gp76 of bacteriophage P23-45 were shown to inhibit *T. thermophilus* transcription (Tagami, Sekine et al. 2014, Ooi, Murayama et al. 2018). Importantly, RNAP is a validated antibacterial target and novel inhibitors for RNAP hold promise for potential new antibiotic development against antimicrobial resistance (Ho, Hudson et al. 2009, Srivastava, Talaue et al. 2011, Ma, Yang et al. 2016).

In this work, we investigate the potential effects of Ocr on the host transcriptional machinery. Our results with purified components show that Ocr can bind to RNAP and inhibit both σ^70^ and σ^54^ dependent transcription. Specifically we establish that an Ocr competes with sigma binding, thus affecting recruitment. Using structural biology, we show that the Ocr dimer directly binds to RNAP at the RNAP cleft, where sigma factors and the transcription bubble bind, suggesting that Ocr interferes with transcription recruitment by competing with sigma factor binding, and also interferes with transcription initiation. Our work thus reveals the detailed molecular mechanisms of how Ocr can influence the host through inhibiting transcription in addition to its role in knocking out restriction and modification. These new structural details of RNAP-Ocr complexes allow comparisons with known transcription initiation complexes and other phage proteins suggesting new avenues to be exploited for inhibiting bacterial transcription and combating antibiotic resistance.

## Results

### Ocr interacts with RNAP with high affinity

An early study by Ratner (Ratner 1974) showed that upon T7 infection, Ocr protein was detected in pull-down experiments using host RNAP as bait, suggesting that Ocr could associate with the host transcriptional machinery. To assess if Ocr can engage with RNAP directly, we carried out *in vitro* interaction studies using purified components (**Figure 1**, **Figure 1 – figure supplement 1**). We observed that purified Ocr co-eluted with RNAP in gel filtration experiments, suggesting a tight interaction of RNAP and Ocr sufficient to withstand the lengthy elution time of the complex during chromatography (**Figure 1A-C**). The complex persisted for the order of minutes during chromatography, suggesting a stable complex. To quantify the interactions, we carried out binding experiments between RNAP and Ocr using microscale thermophoresis (MST). Our results show the binding affinity (dissociation constant) is around 50 nM, similar to that of RNAP with sigma factors **(Figure 1D-F)**. These results thus confirm the basis for the RNAP-Ocr interaction observed by David Ratner many years ago using a pull-down affinity chromatography assay (Ratner 1974). Interestingly, our observed affinity for RNAP is about three orders of magnitude lower than the values observed for the association of Ocr with the EcoKI type I restriction modification enzyme (Atanasiu, Su et al. 2002).

**Figure 1.**
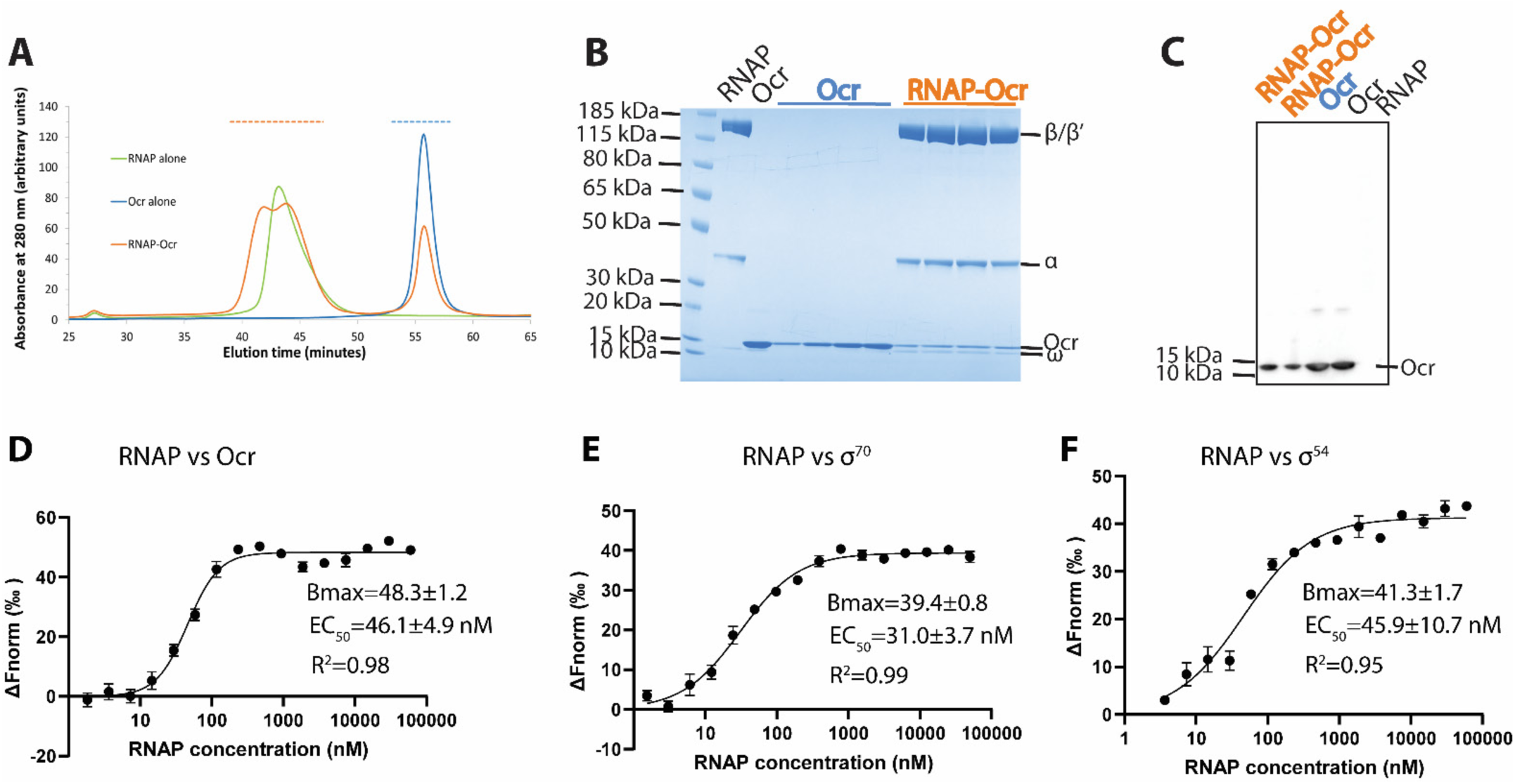
RNAP interacts with Ocr and can form a stable complex with Ocr. **A)** Gel filtration chromatography profiles of RNAP (green), Ocr (blue) and RNAP-Ocr complex (orange). **B)** SDS-PAGE of the corresponding fractions from (A) as indicated by colored dashed lines in (A) and the purified RNAP and Ocr as comparison, the same below in (C). **C)** Western blotting of samples from RNAP-Ocr gel filtration fractions to verify the presence of Ocr co-eluting with RNAP, here only Ocr is his-tagged. **D–F).** MST experiments measure the binding affinity (Kd) of Ocr, σ^70^ and σ^54^ with RNAP.

### Structures of Ocr in complex with RNAP

In order to understand how Ocr interacts with RNAP, we subjected the purified complex from size-exclusion chromatography to cryo electron microscopy (cryoEM) single particle analysis. After several rounds of 2D classification, which allowed us to remove ice contaminations, very large particles or particles much smaller than RNAP, the particles were subject to 3D classifications using the structure of the RNAP filtered to 60 Å as an initial model (PDB code 6GH5, **Figure 2 – figure supplement 1**). Two classes with clear density for Ocr were refined to 3.7 and 3.8 Å resolution respectively (**Figure 2, Figure 2 – figure supplement 2 and supplement 3**). The electron density for RNAP is clear and two distinct structural models of RNAP (taken from PDB code 6GH6 and 6GH5) can be readily fitted into the two reconstructions (**Figure 2 and Figure 2 – figure supplement figure 3A**). The remaining density region that is not accounted for by RNAP can accommodate an Ocr dimer (**Figure 2 – figure supplement 3A**). The density for many side chains of Ocr is visible, which allows accurate positioning of the Ocr atomic structure (PDB code 1S7Z) (**Figure 2 – figure supplement 3B-3C**). The regions with the highest resolution within the reconstruction are in the RNAP core (**Figure 2 – figure supplement 2**), where density for many side chains is clearly visible (**Figure 2 – figure supplement 3C)**.

**Figure 2.**
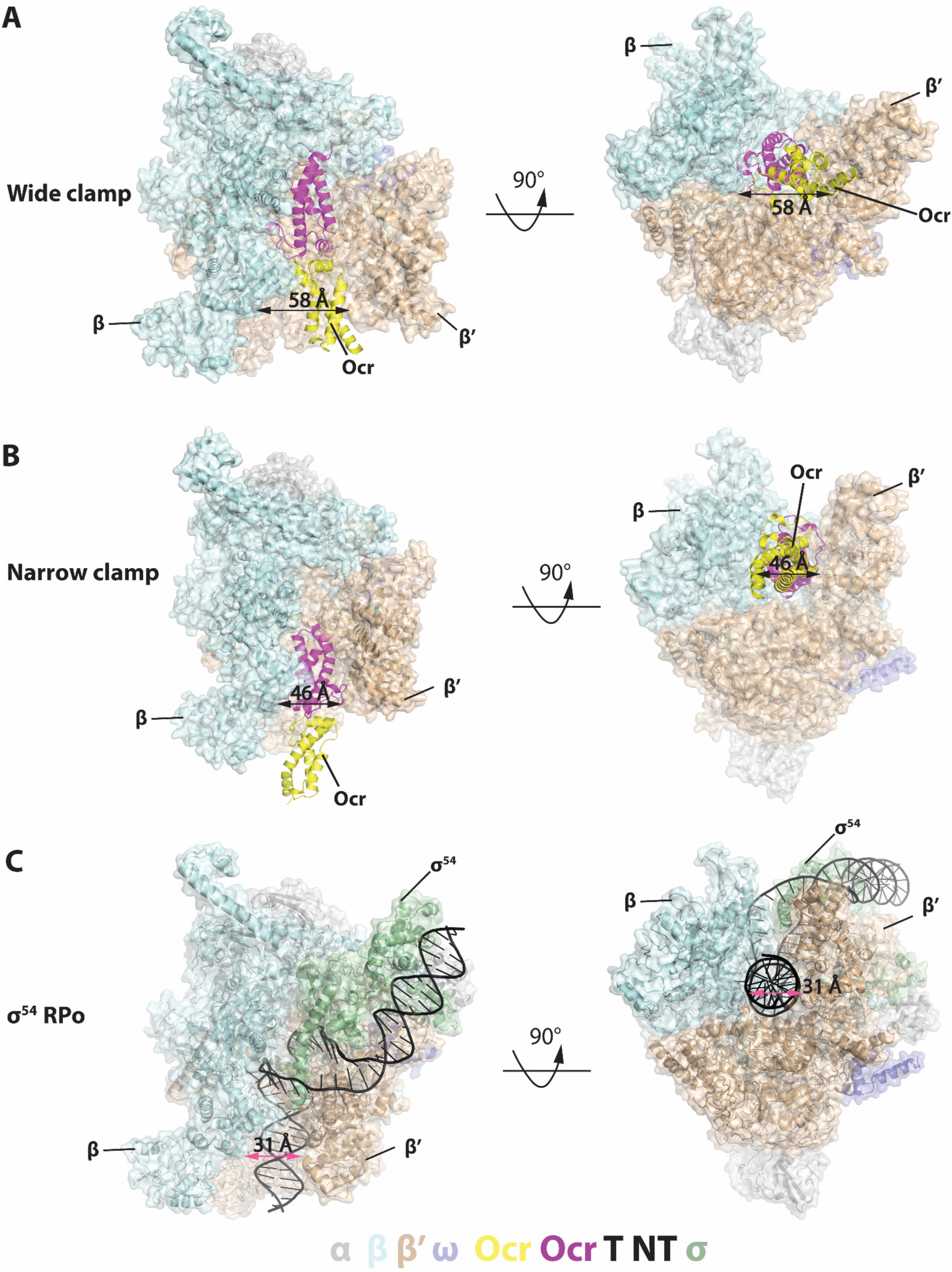
Structures of RNAP-Ocr in two different binding modes in two orthogonal views. **A**) “wide-clamp” mode. **B**) “narrow-clamp” mode. RNAP and σ^54^ were shown as surface, Ocr shows as cartoon. The color key is below the figure. α-grey, β-pale cyan, β’-wheat, ω-slate, σ^54^-palegreen, Ocr–magenta (proximal subunit) and yellow (distal subunit), DNA-black (NT: non-template strand DNA,T: template strand DNA). **C**) RNAP-σ^54^ open promoter complex (PDB code 6GH5).

**Figure 3.**
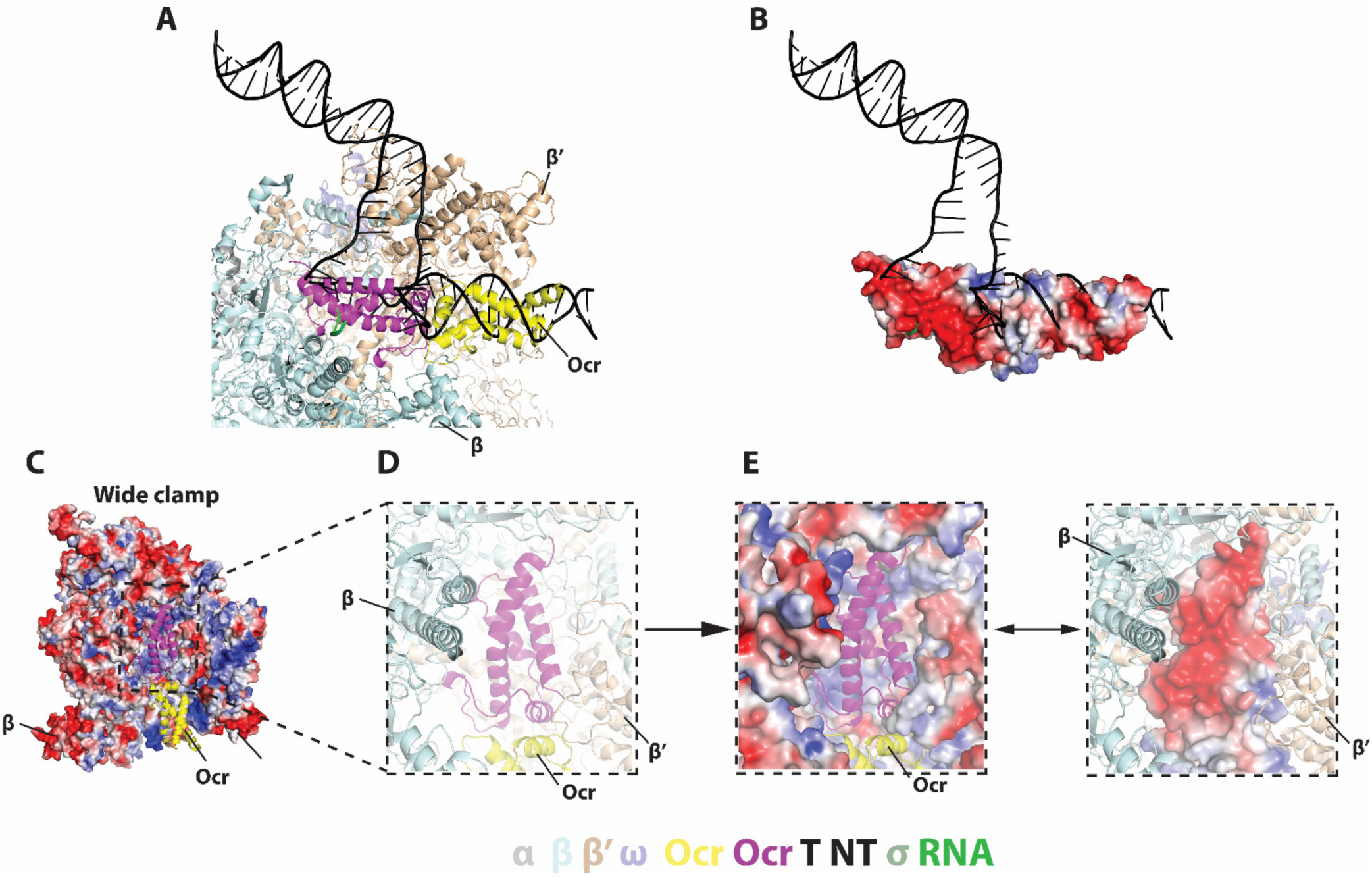
Detailed interactions between Ocr and RNAP and in the “wide-clamp” conformation. **A**)-**B)** comparison with initial transcribing complex structure (PDB 6GFW). **C)** Ocr resides in the positively charged RNAP channel**. D-E**) Detailed charge distributions of RNAP and Ocr in the interacting regions. Blue – positive, red – negative.

During 3D classification, it is clear that a number of different conformational and structural classes exist within the dataset and two distinct classes were identified due to their widely differing RNAP conformations (**Figure 2 – figure supplement 1**). Bacterial RNAP consists of five subunits forming a crab claw shape with the two large subunits β and β’ forming the claws (or clamp) which enclose the RNAP channel that accommodates the transcription bubble and downstream double-stranded (ds) DNA. The RNAP clamp is highly dynamic and single-molecule Forster Resonant Energy Transfer data has shown that the RNAP clamp adapts a range of conformations from closed to open involving more than 20 Å movement and 20° rotation of the clamp (Chakraborty, Wang et al. 2012). Our recent structural work showed that specific clamp conformations are associated with distinct RNAP functional states with the widely opened clamp conformation associated with the DNA loading intermediate state (Glyde, Ye et al. 2018).

In both of the distinct RNAP-Ocr complex structures, Ocr remains as a dimer upon binding to RNAP (**Figure 2A-B**). In one structure, the entire Ocr dimer inserts deeply into the RNAP cleft, occupying the full RNAP nucleic acid binding channel including where the transcription bubble and the downstream DNA resides in the transcriptional open and elongation complexes (**Figure 2A, Figure 2C, Figure 3A**). In this interaction mode, the RNAP clamp is wide open and both Ocr subunits are involved in the interactions (**Figure 2A, Figure 3B, Figure 2 – figure supplement 3**), we thus denote it as the “wide clamp structure”. The second structure involves only one of the two Ocr subunits within the dimer contacting RNAP, occupying only the downstream DNA channel. Here the RNAP clamp is less open and we denote this structure the “narrow clamp structure” (**Figure 2B, Figure 2 – figure supplement 3**).

In the wide clamp structure, the RNAP clamp conformation is very similar to the conformation observed in the transcription initiation intermediate complex where the clamp is wide open and DNA is partially loaded into the cleft (Glyde, Ye et al. 2018)). In this structure, the Ocr dimer inserts into the RNAP channel and each of the two Ocr subunits, which we term proximal and distal, makes distinct interactions with RNAP. The proximal subunit is deeply embedded in the RNAP cleft and occupies the space for the template strand DNA, the complementary newly synthesised RNA and the non-template strand DNA (**Figure 2 and Figure 3, magenta** (Bae, Feklistov et al. 2015, Zuo and Steitz 2015, Glyde, Ye et al. 2018)). The distal Ocr subunit occupies the position of the downstream DNA (**Figure 2 and Figure 3A, yellow**). The negatively charged surface of the proximal Ocr complements the highly positively charged surface of the surrounding RNAP and fits snugly into the channel formed by β and β’ subunits (**Figure 3B-C**). The distal Ocr subunit mimics the downstream DNA in its interactions with the β’ clamp. Superposition of the β’ clamp in the RNAP-Ocr structure with that of open complex structure (RPo) shows the distal Ocr monomer aligns with downstream dsDNA in RPitc (**Figure 3B**). However, the Ocr dimer has a rigid conformation that resembles a bent B-DNA. Consequently, in order to maintain the interactions between both Ocr subunits and RNAP, the β’ clamp has to open up (to ∼ 58 Å in RNAP-Ocr from ∼30 Å in RPo) to accommodate the Ocr dimer compared to RPo that accommodates a transcriptional bubble and dsDNA **(Figures 2 and 3**).

In the narrow clamp mode, the proximal Ocr subunit interacts with the β’ clamp, again mimicking downstream DNA in RPo although the clamp is more open (∼45 Å) compared to that in RPo (∼30 Å) (**Figure 2B-C**). Interestingly, although an Ocr monomer surface mimics that of a dsDNA in the overall dimension and the negative charge distributions, the Ocr subunit in the narrow clamp structure does not exactly overlay with the downstream dsDNA in RPo when RNAP is aligned on the bridge helix, the highly conserved structural feature that is close to the active centre and connects β and β’ clamps (**Figure 4A**). Instead Ocr is shifted upwards compared to the dsDNA in RPo (**Figure 4**). Inspecting the structures of Ocr and RNAP explain the differences. The downstream dsDNA binding channel of RNAP consists of β and β’ clamps on two sides and the β’ jaw domain as a base, providing a positively charged environment on three sides, required to accommodate and engage with the negatively charged DNA during transcription (**Figure 4**). Although an Ocr monomer has a largely negatively charged surface, there are also positively charged areas on the Ocr surface, especially those facing the β’ jaw domain (**Figure 4B right panel**). In order to maintain the interactions between Ocr and the β’ clamp and to overcome charge repulsions with the β’ jaw, the β’ clamp is opened up, shifting Ocr upwards and away from the jaw domain.

**Figure 4.**
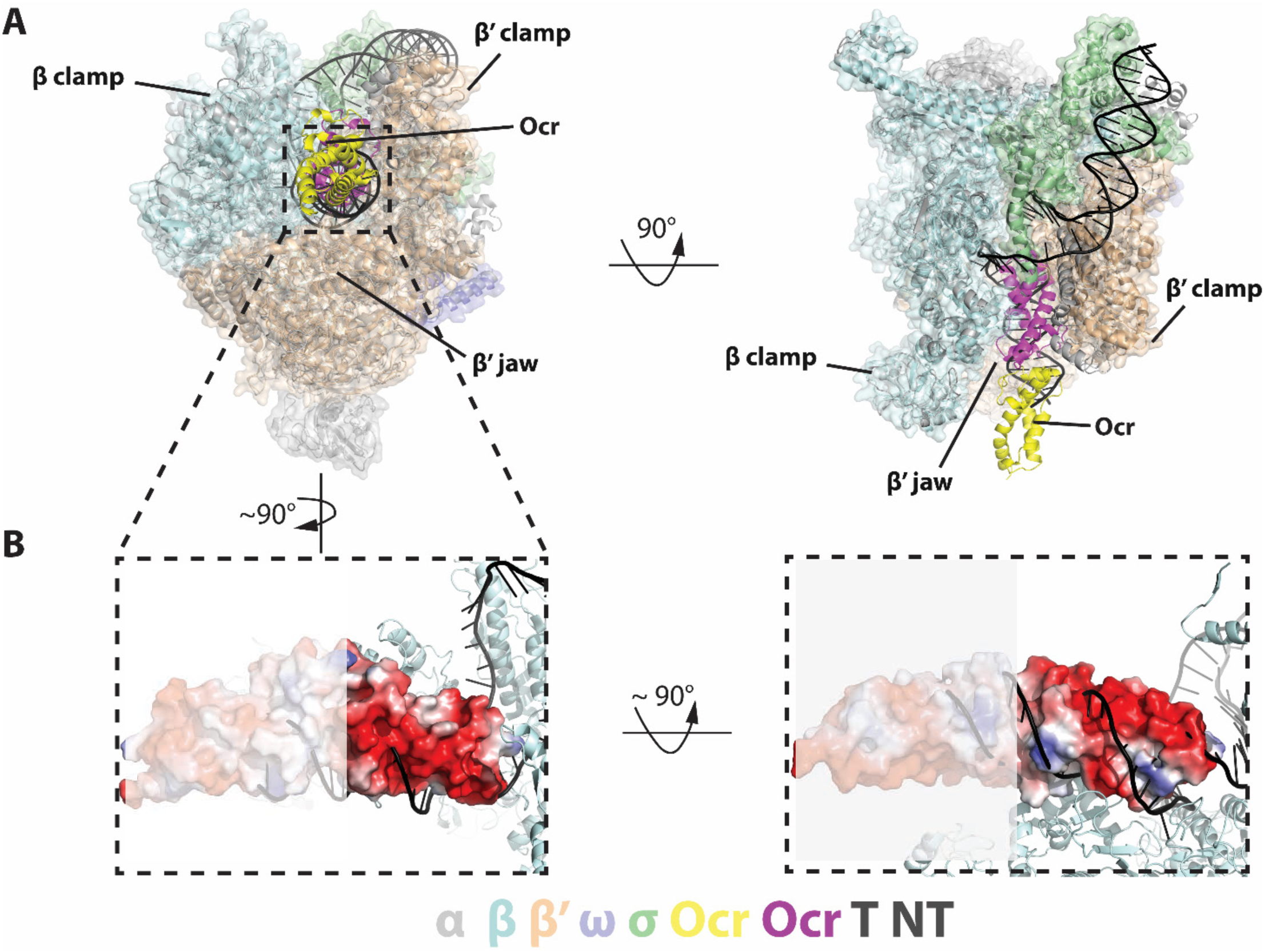
Detailed interactions between RNAP and Ocr in the “narrow-clamp” conformation. **A**) Two orthogonal views of RNAP-Ocr narrow-clamp structure (RNAP shows as surface and Ocr shows as cartoon (magenta and yellow), overlaid with σ^54^ open complex structure. **B**) Insets dash-lined box and its enlarge view shows negative charge distribution of Ocr facing β’ clamp and the orthogonal view shows the slightly positive charge distribution of Ocr facing β’ jaw domain. The distal monomer (yellow) not involved in the interactions are shaded.

### Ocr inhibits transcription in vivo and in vitro

In order to understand the significance of Ocr binding to RNAP, we investigated the effects of Ocr on bacterial transcription. We first analysed the effects of Ocr *in vivo* on cell growth and transcription using *E. coli* MG1655 cells expressing Ocr from a plasmid vector under the control of an arabinose-inducible promoter. The range of growth conditions used are summarised in (**Figure 5 – table supplement 1)**.

**Figure 5.**
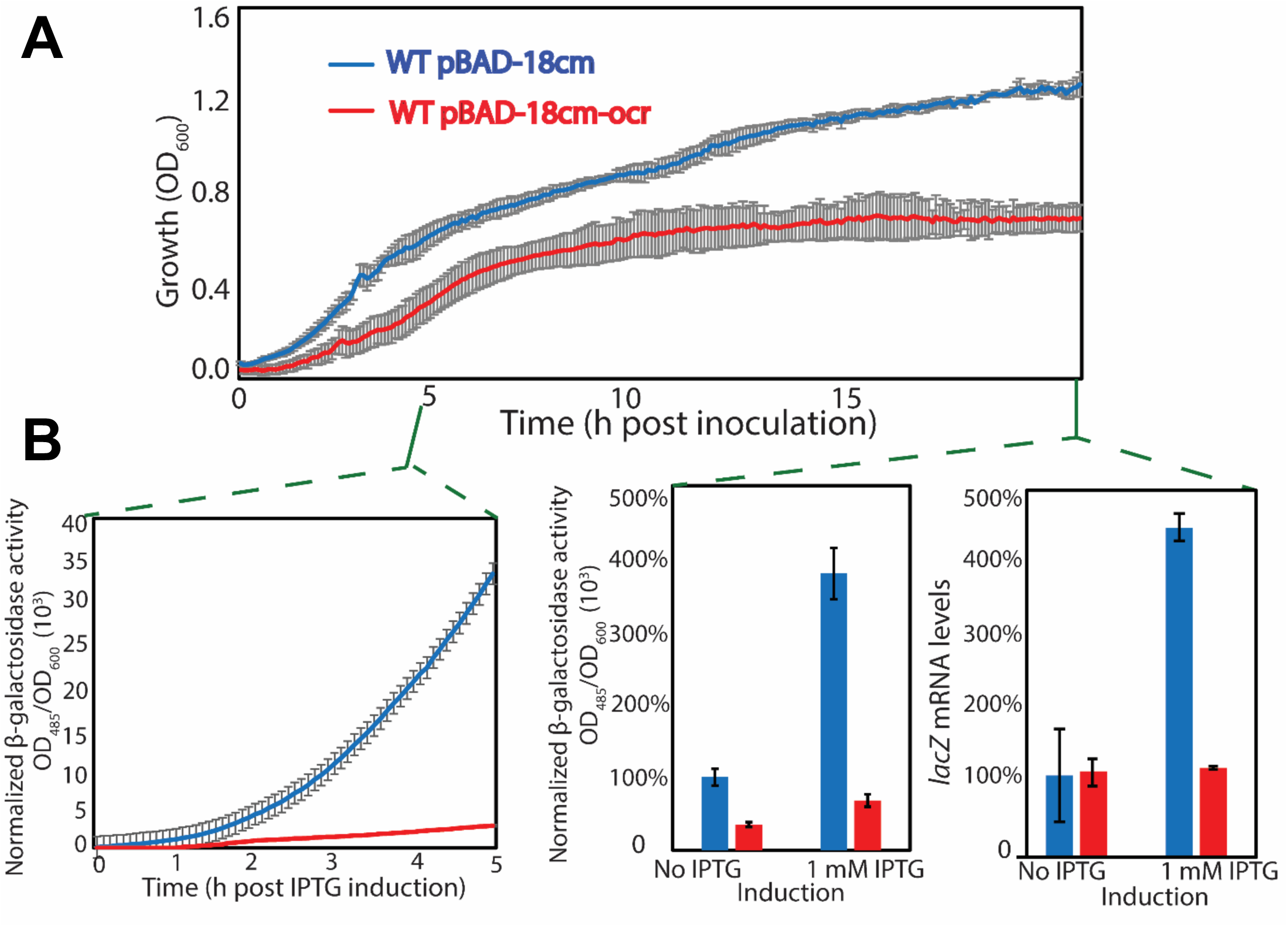
*in vivo* data showing Ocr can inhibit transcription. A) growth of *E. coli* cells in the presence and absence of Ocr in minimal media. B) normalised β-galactosidase activity during a 5 h period (left); β-galactosidase activity (middle) and *lacZ* mRNA levels (right) at 15 h post-induction in the presence and absence of Ocr. At least three independent experiments were performed, each with 3-5 technical replicates.

An Ocr-mediated growth inhibition effect was observed exclusively in minimal modified M9 media, at both temperatures tested, regardless of the level of Ocr inducer (**Figure 5**). The Ocr expression in minimal and complete medium was confirmed using peptide mass fingerprinting by detecting N- and C-terminal tryptic peptides derived from Ocr (**Figure 5 – table supplement 2**). The inhibitory effect of Ocr *in vivo* does not appear to be strongly dependent on inducers, consistent with some leaky expression from the pBAD18cm vector in the absence of arabinose induction. It is possible that addition of extra arabinose does not greatly increase Ocr level because Ocr inhibits its own expression through inactivating RNAP at the promoter that drives Ocr expression. An example growth curve comparison is shown in **Figure 5A**.

The effect of Ocr on transcription in minimal modified M9 medium was further assessed using the chromosomal β-galactosidase gene as a reporter. During the 5 hour period following induction of β-galactosidase with IPTG, a∼10-fold increase of normalised β-galactosidase activity was observed in *E. coli* cells not expressing Ocr compared to those expressing Ocr (**Figure 5B**, left panel). Moreover, at 15 hours post-induction there was a 4-fold increase of normalised β-galactosidase activity in cells not expressing Ocr between induced and un-induced cells, while the β-galactosidase activity in cells expressing Ocr was lower overall and the observed difference between induced and un-induced cells was less than 2-fold (**Figure 5B**, middle panel). Subsequently, measurements of *lacZ* mRNA using reverse transcription followed by quantitative polymerase chain reaction (RT-qPCR) showed that the observed β-galactosidase activity is concomitant with *lacZ* mRNA levels (**Figure 5B**, right panel), confirming that Ocr was inhibitory for transcription *in vivo*.

Having established that Ocr can inhibit transcription *in vivo*, we next assayed its effects *in vitro*. Since Ocr binds to the DNA binding cleft of RNAP, Ocr could potentially interfere with a wide range of transcriptional controls. We thus assessed transcription by the housekeeping σ^70^ as well as the major variant σ^54^ using short-primed RNA (spRNA) assays performed on various promoter DNAs (see Materials and Methods).

To determine the effect of Ocr on RNAP-σ^70^ transcription activity we performed an *in vitro* transcription assay using different promoters and promoter variants such as linear σ^70^ promoter and one variant with a mismatched “pre-opened” DNA sequence (−6 to −1) that favours open complex formation (Zuo and Steitz 2015), or *lac*UV5 promoter (Camara, Liu et al. 2010) as well as a T7A1 supercoiled promoter (**Figure 6 A-C, Figure 6 – figure supplement 1**). Data revealed that adding 5 μM Ocr significantly decreased activity of RNAP-σ^70^ on the different promoters in the range of 90% −30%, while in the presence of equimolar concentrations of Ocr to σ^70^ 0.4 μM Ocr showed a modest inhibitory effect of 30% −10%. On the linear promoter in the presence of 5 μM Ocr the amount of spRNA was significantly reduced when Ocr was added prior or after the holoenzyme formation (**Figure 6A-B** I-II and **Figure 6 – figure supplement 1** I-II) however, the effect of Ocr on RNAP-σ^70^ open complex formation was less pronounced (**Figure 6A-B III** and **Figure 6 – figure supplement 1** III). Ocr did not abolish transcription on the supercoiled T7A1 promoter at the same extent as on the linear promoters, indicating a role for DNA topology and perhaps the energy barriers of DNA opening in open complex formation and promoter sensitivity to Ocr (**Figure 6C** I-III).

**Figure 6.**
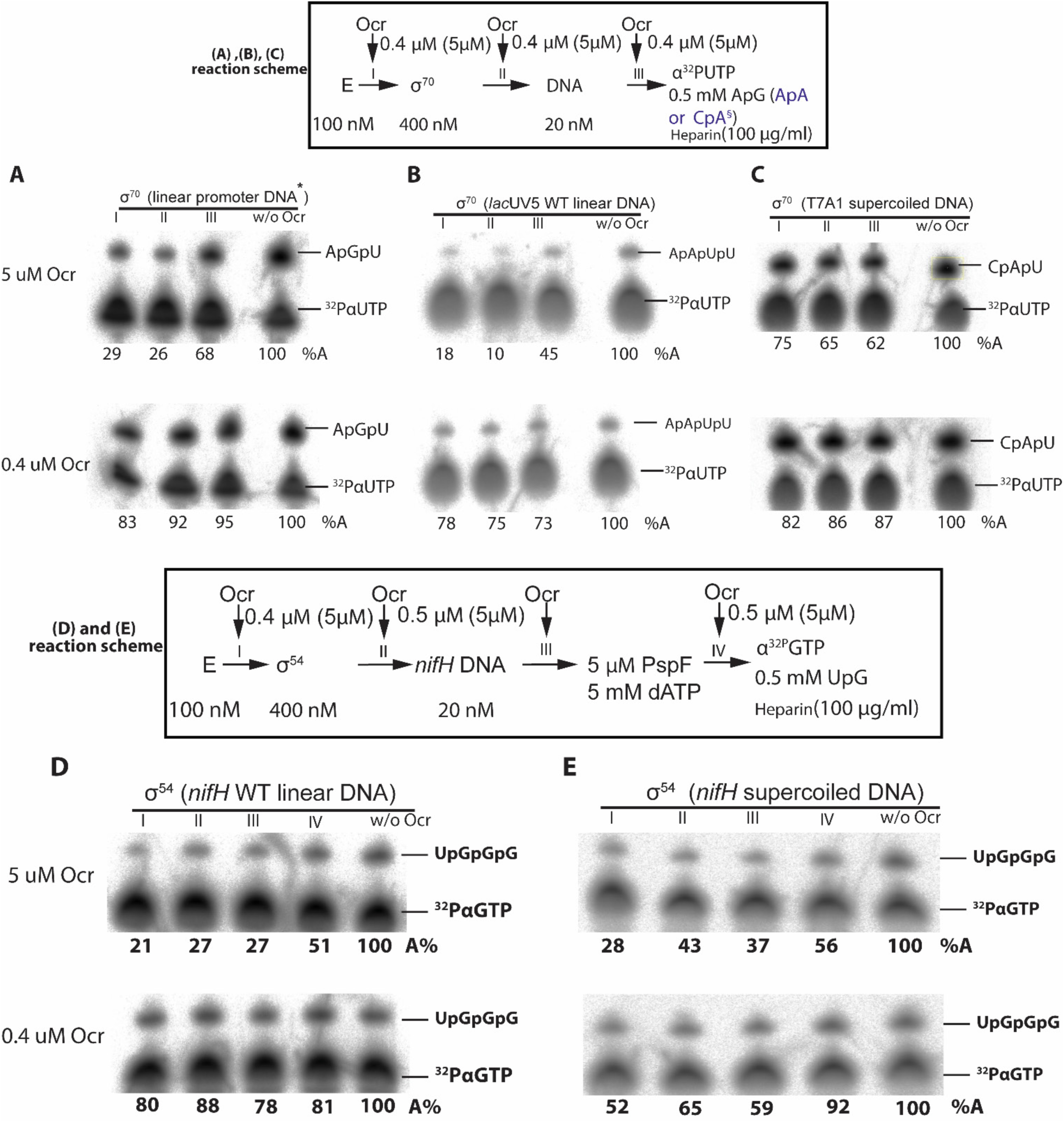
*In vitro* transcription assays. **A)-C**) spRNA experiments on σ^70^ using a range of promoter DNAs. Reaction schematics are shown with I, II, III representing experiments when Ocr is added during different stages transcription initiation. A control lane without Ocr is also shown. **D)-E**) spRNA experiments on σ^54^ using *nifH* linear promoter DNA and supercoil promoter DNA. Reaction schematics are shown with I, II, III, IV representing experiments when Ocr is added during different stages transcription initiation. A control lane without Ocr is also shown. All experiments were performed at least three times and values were within 5% of the relative percentage value shown.

The effect of Ocr on RNAP-σ^54^ dependent transcription was assessed using different variants of *Sinorhizobium meliloti nifH* promoter including linear wild-type, supercoiled wild-type and linear DNA with mismatched bases between −10 and −1 (Chaney and Buck 1999, Glyde, Ye et al. 2018), presenting a pre-opened transcription bubble (**Figure 6 D-E** and **Figure 6 – figure supplement 1**). The PspF_1-275_ protein was used as an activator of RNAP-σ^54^ transcription to promote open complex formation (see Materials and Methods). On both linear and supercoiled *nifH* promoters the RNAP-σ^54^ complexes were sensitive to Ocr, and in the presence of 5μM Ocr transcriptional activity was reduced to 30% of the activity obtained in the absence of Ocr (**Figure 6D-E** I-III). Once the open complex was formed by the action of PspF_1-275_, the Ocr protein showed a 50% inhibitory effect on transcription (**Figure 6D-E** IV). In contrast, the effect of Ocr on the RNAP-σ^54R(336)A^ mutant was lower (**Figure 6 – figure supplement 1**). This mutant was chosen because σ^54^ mutated at R336 to Ala can bypass the requirement of activator protein when the transcription bubble is pre-opened (Chaney and Buck 1999, Xiao, Wigneshweraraj et al. 2009, Glyde, Ye et al. 2018). In the presence of equimolar concentrations of Ocr to σ^54^ (0.4 μM) the inhibitory effect was not as pronounced with around 80% activity on linear *nifH* promoter and 60% activity on supercoiled *nifH* promoter. These data indicate that (i) RNAP core and RNAP-σ^70^ and RNAP-σ^54^ holoenzyme are sensitive to Ocr, (ii) once an open complex is formed the effect of Ocr is reduced and (iii) consistent with (ii) with the RNAP-σ^54R(336)A^ bypass mutant and with pre-opened transcription bubble, the open complex is more efficiently formed than those driven by PspF_1-275_ (**Figure 6E** and **Figure 6 – figure supplement 1B**), thus Ocr is less effective in inhibiting transcription.

### Ocr inhibits RNAP recruitment and open complex formation

Since Ocr interfered with both σ^70^ and σ^54^ dependent transcript formation (above), we wanted to investigate whether it does so through interfering with RNAP recruitment by sigma and/or other processes required for transcription. Specifically, we probed the ability of Ocr in interfering with RNAP-σ holoenzyme formation (recruitment) and the transcription-competent open complex for both σ^54^ and σ^70^.

Using purified components and gel mobility shift assays (**Figure 7**), Ocr was observed to disrupt a pre-formed RNAP-σ^54^ holoenzyme resulting in the formation of an RNAP-Ocr complex. Adding Ocr to pre-formed holoenzyme, the Native PAGE gels clearly showed the gradual disappearance of RNAP-σ^54^ holoenzyme and the appearance of the RNAP-Ocr complex. Due to the lower sensitivity of σ^54^ to coomassie blue staining, the displaced free σ^54^ from the RNAP-σ^54^ holoenzyme was observed by western blotting with an antibody to the His-tag on σ^54^. The σ^54^ was released when equimolar Ocr was added. Increasing Ocr concentration to 4 times that of σ^54^ showed that the pre-formed holoenzyme was almost completely disrupted and much more σ^54^ was released (**Figure 7A** comparing lanes 5-6, 9-12, **Figure 7 – supplement figure 1A**). Furthermore, when Ocr was incubated with RNAP first, the ability of RNAP to form a holoenzyme with σ^54^ was significantly reduced (comparing **Figure 7A** lanes 5-8). These results thus demonstrate that Ocr inhibits and disrupts the formation of RNAP-σ^54^ holoenzyme *in vitro*. Similar experiments with σ^70^ showed that Ocr prevents RNAP-σ^70^ holoenzyme formation and disrupts pre-formed RNAP-σ^70^ holoenzyme. The MST experiment of Ocr binding to RNAP-σ^70^ revealed that Ocr can displace σ^70^ with an IC50 of 41 nM (**Figure 7B, figure 7 – supplement figure 1B-C**).

**Figure 7.**
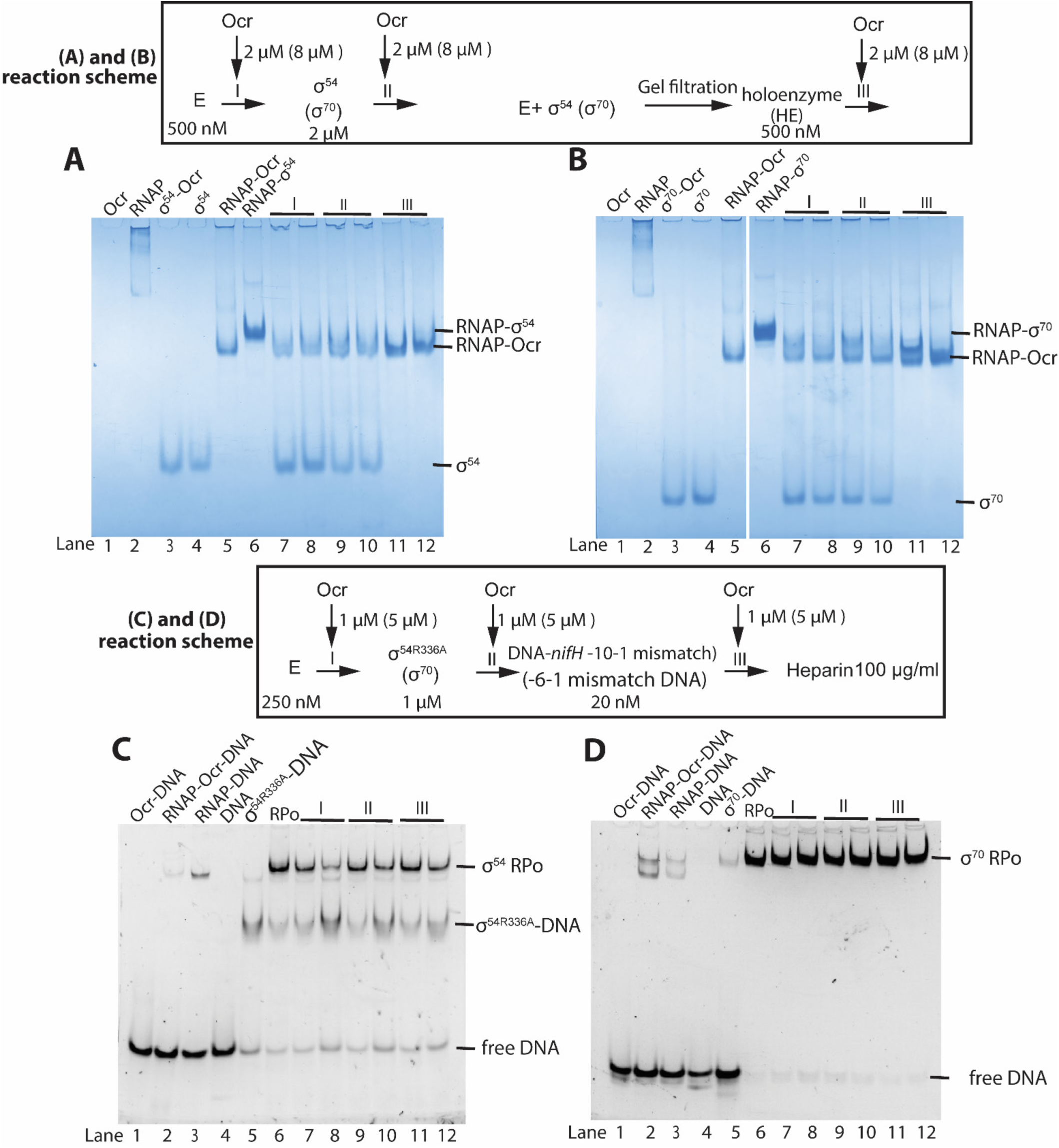
Competition experiments of Ocr on holoenzyme and open complex formation of σ^54^ and σ^70^ as assayed by native-PAGE gels. **A**) Ocr and σ^54^ holoenzyme formation, **B**) Ocr and σ^70^ holoenzyme formation. Reaction scheme are indicated above. E indicates core enzyme RNAP, I, II, III indicate the point when Ocr is added. Protein and DNA concentration are shown. In lane 7, 9 and 11, a 1:1 molar ratio of Ocr to σ was used, whereas in lanes 8, 10, 12, a 4:1 ratio of Ocr to σ was used. **C**) Ocr and its effect on σ^54^ open complex formation, **D**) Ocr and its effect on σ^70^ open complex formation, The reaction schematics are shown, I, II, III indicates that point when Ocr was added during the reaction. For open complex in (C) and (D), all the reactions including 100 µg/ml heparin.

The open complex represents a transcription-ready functional complex. Using promoter DNA containing pre-opened transcription bubbles allows formation of stable open complexes for RNAP-σ^70^ and RNAP-σ^54R336A^. Interestingly, preincubation of RNAP with excess of Ocr showed that the ability of RNAP to form open complex with σ^54^-DNA or σ^70^-DNA was less pronounced (**Figure 7D-E I**). The inhibition is less effective compared to its effects on holoenzyme (**Figure 7A-B** lanes 7-10, I, II). Once the open complex is formed, the ability of Ocr to disrupt the open complex is limited (compare **Figure 7D-E** III), suggesting that Ocr mainly acts at the initial recruitment stage and open complex formation, consistent with the *in vitro* transcription results (**Figure 6**). However, given the limited effects on pre-opened open complex, Ocr is unlikely to act effectively on an actively transcribing RNAP.

## Discussion

### Ocr is a bifunctional protein

Our data here demonstrate a new activity for Ocr in inhibiting the host transcriptional machinery in addition to its role in inhibiting the type I restriction/modification systems of the host. Thus Ocr is a bifunctional DNA mimic protein.

Structures of Ocr in complex with RNAP and the *in vitro* Ocr competition experiments clearly show that Ocr can inhibit transcription. The structures of Ocr-RNAP show that Ocr occupies the space where RII.1-RII.3 of σ^54^ and σ_1.1_ and σ_3.2_ of σ^70^ reside, thus it would directly compete with σ^70^ and σ^54^ binding, inhibiting holoenzyme formation (**Figure 8A-B** (Murakami 2013, Yang, Darbari et al. 2015)). Even when holoenzyme is pre-formed, Ocr can compete with σ for RNAP binding. The cleft where Ocr binds is also where the transcription bubble and downstream DNA reside in the open complex and elongation complex, thus explaining how Ocr inhibits open complex formation (Vassylyev, Vassylyeva et al. 2007, Zuo and Steitz 2015).

**Figure 8.**
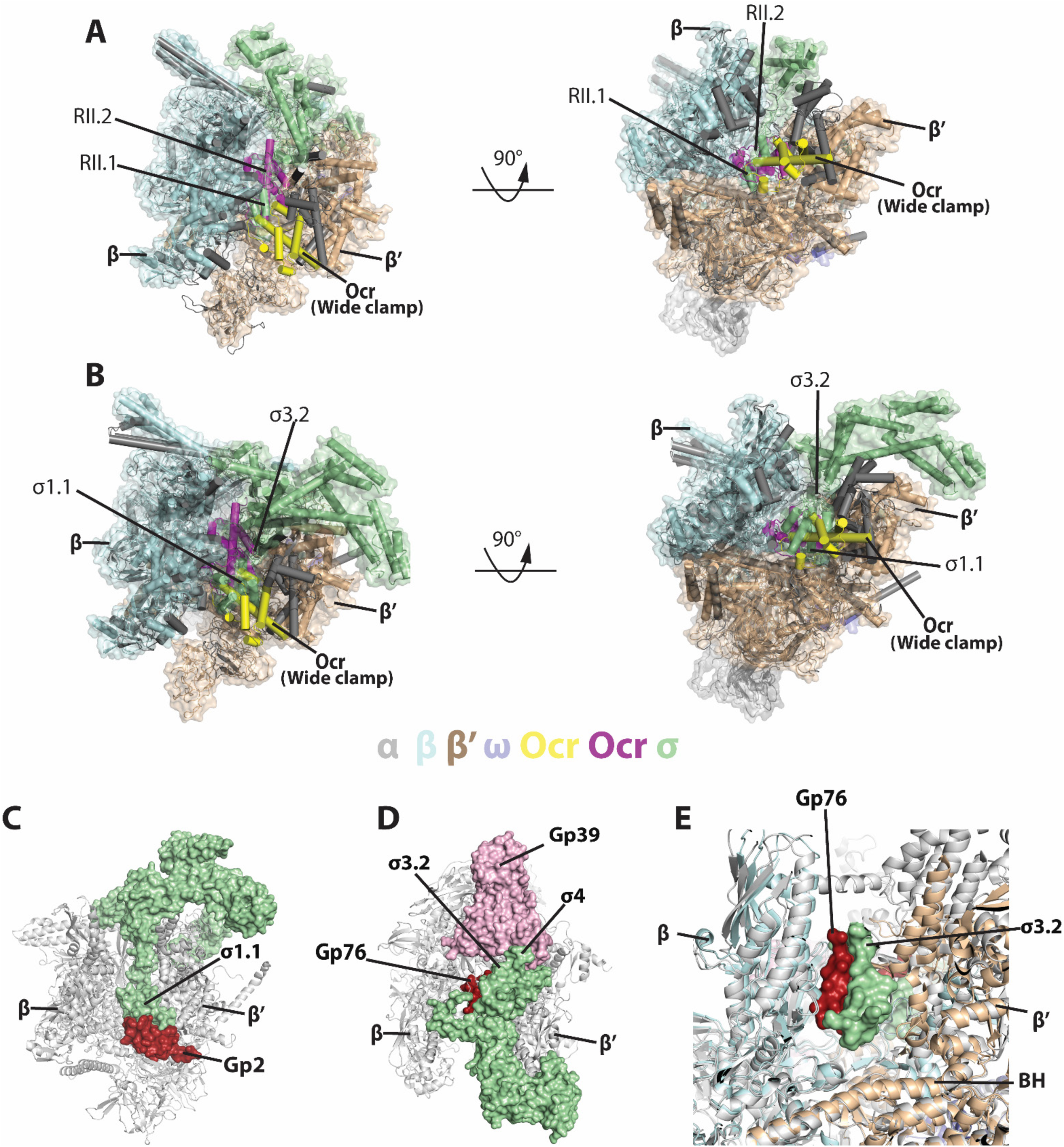
Comparisons with sigma and other phage proteins in binding to RNAP. **A**) Ocr overlay with RNAP-σ^54^ holoenzyme, **B**) Ocr overlay with RNAP-σ^70^ holoenzyme **C**) Complex structure of gp2 of T7 with σ^70^ holoenzyme (PDB 4LLG), **D**) Complex structure of gp76-gp39 of P23-45 with σ^70^ holoenzyme (PDB 5XJ0), **E**) Enlarged view of gp76 and σ^70^. RNAP shows as cartoon and coloured as grey, σ^70^ shows as surface and coloured pale green. Gp2, gp39, gp76 are show as surface and coloured as firebrick, light pink and firebrick, respectively.

*In vitro* transcription results show a larger effect when Ocr is added prior to holoenzyme formation but only limited effects once the open complex forms. This dependence on the stage at which Ocr is added to RNAP is reminiscent of the dependence on the stage at which Ocr is added to a Type I restriction/modification reaction (Bandyopadhyay, Studier et al. 1985). These results are in agreement with direct competition experiments (**Figure 7**) showing that Ocr effectively inhibits holoenzyme formation but has only limited effects on the pre-formed open complex and or transcribing RNAP. This is consistent with the very slight effects on transcription in rich media, as under rapid growth conditions where transcriptional activities are high, the majority of the RNAP in cells is associated with DNA and is engaged in transcription, either as open promoter complexes or elongation complexes. However, using minimal media where growth is slowed down, transcription activities are reduced. Here some RNAP molecules presumably are not associated with DNA allowing Ocr to effectively inhibit holoenzyme and open complex formation, explaining its growth conditional inhibitory effect *in vivo (***Figure 5**).

Our data suggest an answer to the puzzle of why Ocr is so abundantly overexpressed immediately upon a T7 infection (Issinger and Hausmann 1972), when its classical target, the Type I restriction/modification enzyme, is present in very low numbers (estimated at ∼60 molecules per cell, (Kelleher and Raleigh 1994)). The very tight binding between Ocr and Type I restriction/modification enzymes ensures complete inhibition of restriction and modification. It would appear from our data that the excess Ocr molecules could start the process of shutting down host transcription by binding to any non-transcribing RNAP. Subsequently the expression of other phage proteins, gp0.7, gp2 and gp5.7, completes the shutdown of host RNAP. This staged but possibly coordinated inhibition of host machinery might play an important role in phage infection and requires further *in vivo* investigations.

The bifunctionality observed in Ocr may also be shared by other DNA mimic proteins. For example, the phage T4 Arn protein targeting the McrBC restriction enzyme also has weak interaction with histone-like protein H-NS (Ho, Wang et al. 2014). Arn is very similar in shape to Ocr but since T4 requires the host RNAP throughout its life cycle it is unlikely to target RNAP as strongly as Ocr. Additional functions may also exist for other DNA mimics like phage λ Gam protein (Wilkinson, Troman et al. 2016) and the recently discovered anti-CRISPR proteins (Wang, Chou et al. 2018). How the multifunctionality of these proteins are utilised by phage is an interesting topic for further study.

### Comparisons with other RNAP inhibiting bacteriophage proteins

Several other bacteriophage proteins have been reported to inhibit transcription. These include T7 gp2 and gp5.7, which inhibits *E. coli* RNAP, and P23-45 gp39 and gp76, which inhibit *T. thermophilus* transcription. Structural and biochemical studies show that gp2 specifically inhibits σ^70^-dependent transcription through the interactions and stabilisation of the position of the inhibitory region 1.1 of σ^70^, between the β and β’ clamps at the rim of the downstream DNA binding channel (**Figure 8C**, (Bae, Davis et al. 2013). Gp39 binds at region 4 of σ^A^ and restricts its conformation so that it is unable to bind to −35 promoter regions (**Figure 8D**, (Tagami, Sekine et al. 2014)). Gp76 on the other hand, binds deeply in the RNAP cleft, preventing open complex formation as well as transcription elongation (**Figure 8D**, (Ooi, Murayama et al. 2018)). However, the binding site of gp76 is adjacent to, but not overlapping the σ^70^ region σ_3.2_, thus gp76 does not inhibit holoenzyme formation (**Figure 8E**).

In all the cases above, the bacteriophage proteins do not inhibit holoenzyme formation. Ocr on the other hand, inhibits holoenzyme formation *per se* and can disrupt a pre-formed holoenzyme (**Figure 7A-B**). Structural comparisons show that Ocr occupies spaces deep inside the RNAP cleft, where normally σ^70^ region 1.1 and region 3.2, as well as σ^54^ region II reside. Ocr binding would thus prevent σ binding as well as destabilise bound σ, preventing holoenzyme formation. Ocr can also inhibit open complex formation but is unable to effectively disrupt pre-formed open complex (**Figure 7C-D** III). This is due to the stable nature of open complex and the overlapping binding sites of Ocr and DNA to RNAP. Furthermore, the clamp is closed in the open complex, a conformation incompatible with Ocr binding (**Figure 2**).

### Unique features of Ocr defines its ability in specific DNA processing systems

Ocr was shown to mimic a dsDNA in both shape and charge distribution (Walkinshaw, Taylor et al. 2002) and an Ocr dimer mimics a slightly bent B-form DNA. Ocr has been shown to occupy the DNA binding grooves of type I restriction/modification enzymes through specific interactions, thus preventing viral DNA from being degraded or modified by the host restriction/modification system (Atanasiu, Su et al. 2002, Kennaway, Obarska-Kosinska et al. 2009).

The structural model of the archetypal Type I restriction/modification enzyme EcoKI (Kennaway, Obarska-Kosinska et al. 2009) shows a close association of Ocr with the M.EcoKI modification methyltransferase core of the Type I restriction/modification enzyme (Kennaway, Taylor et al. 2012). The surface area of the methyltransferase is 76,152 Å^2^ and that of the complex is 84,055 Å^2^. The surface area of the Ocr dimer (Walkinshaw, Taylor et al. 2002) (PDB 1S7Z) is 13,130 Å^2^, thus 2614 Å^2^ is buried on the interface of Ocr with M.EcoKI. The surface area buried in the wide and narrow complexes of Ocr with RNAP is 2573 Å^2^ and 1811 Å^2^ respectively. Therefore it is not surprising that both Ocr-RNAP and Ocr-M.EcoKI complexes are sufficiently stable to persist during lengthy size exclusion chromatography experiments (**Figure 1** and (Atanasiu, Su et al. 2002)).

In this work, we show that Ocr remains as a dimer and binds at the downstream DNA channel of RNAP via the complementary charge interactions between the positively charged RNAP channel and the negatively charged Ocr dimer. However, due to the extensive positively charged surface within RNAP, the binding is not specific. Indeed, two distinct binding modes have been detected which allow either the proximal or distal Ocr monomer to interact with the positively charged β’ clamp. The RNAP clamp opens up in the “wide clamp” binding mode in order to accommodate the Ocr dimer conformation while maintaining the charge complementarity, both deep inside the cleft and with the β’ clamp. When the clamp is closed as in the transcriptional open complex, even though there is ample space to accommodate one Ocr subunit, the charge repulsion between the β’ jaw domain and the Ocr surface results in the opening of the β’ clamp, thus shifting Ocr upwards and away from the β’ jaw domain. These structures demonstrate that the structural flexibility of RNAP clamp plays a key roles in its ability to accommodate Ocr. A study of the kinetics of the formation of the complexes between Ocr and RNAP would be interesting and may show multiple consecutive reactions as found for the interaction of a small DNA mimic Uracil glycosylase inhibitor (UGI) with uracil glycosylase (Bennett, Schimerlik et al. 1993).

The interactions and structures of Ocr with RNAP observed here suggest that Ocr could potentially interact and inhibit other DNA processing enzymes through a series of non-specific interactions with dsDNA binding sites/channels. However, the rigidity and the bent conformation of the Ocr dimer imposes constraints on the binding sites, which are optimised for the restriction enzymes it inhibits but can still be accommodated by the structurally flexible RNAP DNA binding channel. The rigidity of the dimer is defined by the interactions between the Ocr subunits. Work presented here suggests that Ocr could be modified to fine-tune its DNA mimicry for binding to specific DNA processing proteins, either by reducing the rigidity of the dimer or changing the bend angle to create a new dimer interface. DNA mimicry has been proposed as a potential effective therapeutic tool (Putnam and Tainer 2005, Roberts, Stephanou et al. 2012), Ocr could thus be exploited to create specific variants that can target specific DNA binding proteins, especially those that are shown to be antibiotic targets. In addition, our data here show that Ocr can effectively inhibit transcription under stress conditions when transcriptional activity is reduced. It is thus possible that during other growth conditions/phases, such as stationary phases or persistence, Ocr could be effectively utilised to inhibit transcription and thus bacterial survival.

## Materials and Methods

### RNAP-Ocr complex formation

*E. coli* RNA polymerase was purified as previously reported (Yang, Darbari et al. 2015). As well as being purified as previously described (Sturrock 2001), Ocr was ligated into pOPINF vector with a N-terminal His-tag. Three purification steps (Ni-NTA affinity, Heparin, and Gel filtration chromatography) were employed, which generated homogeneous Ocr protein as judged by SDS-PAGE gel. The RNAP-Ocr complex was purified by gel filtration chromatography. Initially RNAP (final concentration 14.5 µM) and Ocr (all Ocr concentrations are for the dimeric form, final concentration 58.0 µM) were mixed at a 1:4 molar ratio in a 150 µl reaction volume and the mixture (150 µl) was incubated on ice for 30 mins before being loaded onto the gel filtration column with a flowrate of 0.3 ml/min (Superose 6 10/300, GE Healthcare) and eluted in buffer containing 20 mM Tris pH8.0, 150 mM NaCl, 1 mM TCEP, 5 mM MgCl_2_. The fractions were run on SDS-PAGE and the presence of Ocr was confirmed by Western blot against the His-tag. The same amount and concentration of RNAP and Ocr alone were also loaded on a Superose 6 10/300 column separately as a control subsequently.

### Microscale thermophoresis (MST)

All MST experiments were performed using a Monolith NT.115 instrument (NanoTemper Technologies, Germany) at 22 ℃. PBS (137 mM NaCl, 2.7 mM KCl, 10 mM Na_2_HPO_4_, 1.8 mM KH_2_PO_4_) supplemented with 0.05% Tween-20 was used as MST-binding buffer for all experiments. In all cases hydrophobic treated capillaries were used. σ^54^ and σ^70^ were purified to homogeneity as described previously (Nechaev and Severinov 1999, Yang, Darbari et al. 2015). Ocr, σ^54^, and σ^70^ were all His tagged and labelled with the kit (Monolith His-Tag Labeling Kit RED-tris-NTA 2nd Generation) separately. The labelled Ocr and sigma factors were diluted to a concentration of 50 nM and mixed with an equal volume of a serial dilution series of RNAP incubated at room temperature for 20 min before loading into MST capillaries. For Ocr competitive binding with σ^70^ holoenzyme, 50 nM of his-labeled σ^70^ holoenzyme, diluted in MST-binding buffer, was mixed with equal volumes of a dilution series of Ocr in MST-binding buffer. Single MST experiments were performed using 70% LED power and 40-60% MST power with a wait time of 5 s, laser on time of 30 s and a back-diffusion time of 5 s. MST data were analyzed in GraphPad Prism and the data were fitted with the Hill equation. The mean half effective concentration (EC50) values were calculated with standard error (SE). In the competitive binding experiment with Ocr and σ^70^-RNAP holoenzyme, the data were fitted using on a variable slope model [log(inhibitor) vs. response curves] from which one could determine the IC50 of Ocr [the concentration of that provokes a response half way between the maximal (Top) response and the maximally inhibited (Bottom) response). Each experiment was repeated at least three times.

### CryoEM Sample preparation

For structural studies using cryoEM, the His-tag of Ocr was cleaved and the purification procedure was as previously reported (Sturrock, Dryden et al. 2001). The complex was formed as described above. The RNAP-Ocr complex sample was concentrated to 0.6 mg/ml and 3.5 µl of the complex was applied to R2/2 holey carbon grids (Quantifoil). The vitrification process was performed using a Vitrobot Mark IV (FEI) at 4 ℃ and 95% humidity, blotted for 1.5 s with blotting force −6. The grid was then flash frozen in liquid ethane and stored in the liquid nitrogen before data collection.

### Electron microscopy data collection

The cryoEM data were collected at eBIC (Diamond Light Source, UK) on a Titan Krios using EPU (FEI) operated at 300 kV and a K2 summit direct electron detector (Gatan). The data were collected with a defocus (underfocus) range between 1.5 µm to 3.5 µm. A total of 3543 micrographs were collected at pixel size of 1.06 Å/pixel and a dose of 50 e^−^/Å^2^ and each micrograph was fractioned into 41 frames (1.21 e^−^/Å^2^/frame).

### Image processing

The procedure of data processing is summarised in **Fig. S2**. Frame alignment and dose weighting were carried out with MotionCor2 (Zheng, Palovcak et al. 2017) before estimating CTF parameters using Gctf (Zhang 2016) and particle picking using Gautomatch (https://www.mrc-lmb.cam.ac.uk/kzhang/Gautomatch/) without a template. The picked particles were then extracted into boxes of 256 × 256 pixels. Initial 2D classification of the data was carried out in Cryosparc (Punjani, Rubinstein et al. 2017) to remove junk particles due to ice contamination or other defects on the grids. Subsequent image processing was carried out in Relion 2.1 (Scheres 2012). Briefly, the particles were separated using 3D classification. Three out of five classes were subsequently refined using the RNAP from the closed complex RPc (EMD-3695) filtered to 60 Å as the initial reference map. The remaining two classes were discarded due to their lack of clear structural features, probably derived from particles that are of poor quality. After combining the remaining three classes and refinement, one more round of focused 3D classification by applying a mask around Ocr was carried out to separate different complexes or conformations. Two classes with clearly different conformations of RNAP and both with Ocr bound, were refined, polished and post-processed (with masking and sharpening), resulting in the final reconstructions at 3.7 Å (for the wide clamp RNAP-Ocr) and 3.8 Å (for the narrow clamp RNAP-Ocr) based on the gold-standard Fourier shell correlation (FSC) at 0.143 criterion.

### Model building, refinement and structural analysis

The RNAP from RPo (PDB code: 6GH5) and RPip (PDB code: 6GH6) was used as an initial model for the model building of narrow clamp and wide clamp reconstructions, respectively. Briefly, the RNAP was first fitted into the RNAP-Ocr density map in Chimera (Goddard, Huang et al. 2007). Subsequently the RNAP structure was subject to flexible fitting using MDFF (Trabuco, Villa et al. 2009). The Ocr crystal structure (PDB code: 1S7Z) was manually fitted into the extra density of the RNAP-Ocr map in Coot (Emsley, Lohkamp et al. 2010). Jelly body refinement in Refmac (Murshudov, Skubak et al. 2011) and real space refinement in Phenix (Afonine, Grosse-Kunstleve et al. 2012) were used to improve the model quality. The final statistics of the model are in **Table 1**. The figures used for structure analysis and comparison were produced in Pymol (The PyMOL Molecular Graphics System, Version 2.0 Schrödinger, LLC) and UCSF Chimera (Goddard, Huang et al. 2007).

**Table 1.** Cryo-EM data collection and refinement statistics

|  |  |  |
| --- | --- | --- |
| **Data collection and Processing** |  |  |
| Magnification | 47393 |  |
| Total micrographs | 3543 |  |
| Movie frames | 41 |  |
| Pixel size (Å) | 1.055 |  |
| Defocus range (µm) | -1.2 to -3 |  |
| Voltage (kV) | 300 |  |
| Electron dose (e<sup>-</sup>/Å<sup>-2</sup>) | 49.53 |  |
| Total particles | 753783 |  |
| FSC threshold | 0.143 |  |
| **3D Reconstruction(RELION)** | **Wide clamp** | **Narrow clamp** |
| Particles | 33646 | 27312 |
| Resolution (Å) | 3.7 | 3.8 |
| **Model refinement and quality** |  |  |
| Resolution (Å) | 3.7 | 3.8 |
| **R.m.s.deviation** |  |  |
| Bond length (Å) | 0.003 | 0.003 |
| Bond angle (°) | 0.788 | 0.813 |
| **Ramachandran plot** |  |  |
| Favored regions (%) | 92.03 | 91.43 |
| Allowed regions (%) | 7.80 | 8.48 |
| Outlier | 0.18 | 0.09 |
| **Validation** |  |  |
| All-atom clashscore | 4.16 | 4.17 |
| Rotamer outliers (%) | 0.19 | 0 |
| C-beta deviations | 0 | 0 |

### Native gel mobility assay and western blotting

σ^54^ and its mutant σ^54R336A^ as well as σ^70^ were purified to homogeneity as described previously (Nechaev and Severinov 1999, Yang, Darbari et al. 2015). All the reactions were carried out in the binding buffer (20 mM Tris-HCl pH8.0, 200 mM KCl, 5 mM MgCl_2,_ 1 mM DTT, 5% glycerol) at room temperature, for the sequentially addition of different components in each step, the incubation time was approximately15-20 min and all the gels used were polyacrylamide. To test the effects of Ocr on holoenzyme formation for both σ^54^ and σ^70^, all the reactions were carried out in 20 µl at room temperature. The order of addition, final concentrations and molar ratio of different components are listed in corresponding figures. For each 20 µl reaction, 10 ul was taken out and loaded into one polyacrylamide gel (4.5%) and the remaining 10 ul of each sample was loaded to another identical gel (4.5%) and run in parallel for 160 mins at 80 volts at 4 ℃ cold room. One gel was then stained with Coomassie blue and visualised, whilst the other gel was used for western blotting against the His-tagged protein of either σ^54^ or σ^70^, following the standard western blotting protocol (iBlot 2 Dry Blotting System, Thermo Fisher Scientific).

To test the effects of Ocr on the formation of the σ^54^ and σ^70^ open complexes, DNA probes used in this study were all Cy3 labelled and the promoter sequences used for σ^54^ open complexes is the same as in our previous work (Glyde, Ye et al. 2017, Glyde, Ye et al. 2018). For σ^70^ open complex experiments, the non-template DNA used was 5’-ACTTGACATCCCACCTCACGTATGCTATAATGTGTGCAGTCTGACGCGG-3’, and the template DNA used was 5’-CCGCGTCAGACTCGTAGGATTATAGCATACGTGAGGTGGGATGTCAAGT-3’. All the other reactions were carried out at room temperature. The sequence of addition, the final concentration, molar ratios of different reactions are described in respective figures. All the gel (5 %) are run at 4℃in cold rooms, with 100 volts for 75 mins. The results were analysed by visualizing and quantifying the fluorescent signals of Cy3, attached to the DNA probes.

All the experiments were repeated at least once and the results are consistent with each other.

### Small primed RNA assays

All reactions were performed in 10 ul final volumes containing: STA buffer (Burrows et al., 2010), 100 nM holoenzyme (1:4 ratio of RNAP:σ^54^), 20 nM promoter DNA probe (for σ^54^ dependent transcription open complex formation, 4 mM dATP and 5 mM PspF_1-275_ were also present) and incubated at room temperature. The sp RNA synthesis was initiated by adding 0.5 mM dinucleotide primer (ApG, ApA, CpA or UpG), 0.2 mCi/ml [α-^32^P] GTP (3000 Ci/mmol) or 0.2 mCi/ml [α-^32^P] UTP (3000 Ci/mmol). The reaction mixtures were quenched by addition of 4 µl of denaturing formamide loading buffer and run on a 20% denaturing gel and visualised using a Fuji FLA-5000 Phosphorimager. At least three independent experiments were carried out and values were within 5% of the relative % values measured.

### Bacterial growth curves

Growth of *E. coli* MG1655 cells not expressing Ocr (WT/pBAD18cm) and expressing Ocr (WT/pBAD18cm[*ocr*]) was monitored for 20 hrs using a FLUOstar Omega plate reader (BMG LABTECH). Starting cultures of 0.1 OD_600_ were inoculated in four different media, Luria-Bertani Broth (LB, Bertani REF) with low (5 g/L) and high (10 g/L) salt concentration, nutrient broth (Oxoid) and modified M9 medium (Teknova), at two different temperatures, 37°C and 25°C. Expression of Ocr was induced using two different concentration of arabinose, 0.02%(^w^/_v_) and 0.2%(^w^/_v_).

### Beta-galactosidase assays

Gene expression levels of beta-galactosidase in *E. coli* MG1655 cells not expressing Ocr (WT/pBAD18cm) and expressing Ocr (WT/pBAD18cm[*ocr*]) were assessed using 0.002 mg/ml fluorescein di(beta-D-galactopyranoside) (FDG; Sigma-Aldrich). FDG results in fluorescence signals, which are proportional to the enzymatic activities after being hydrolysed to fluorescein by beta-galactosidase. Beta-galactosidase expression was induced by 1 mM IPTG 5 hrs post-inoculation.

### RNA extraction, DNase digestion, reverse transcription and quantitative polymerase chain reaction (RT-qPCR)

Following the beta-galactosidase assays, total bacterial RNA was extracted from *E. coli* MG1655 cells using the RNeasy Mini Kit (Qiagen) according to the manufacturer’s instructions. Residual DNA was digested by DNase I (Promega) and cDNA synthesis was performed using 500 ng of RNA and the SuperScript IV reverse transcriptase (Invitrogen) according to the manufacturer’s instructions. Quantitation of the beta-galactosidase mRNA levels was performed using the specific oligonucleotide primers 5’-ATG GGT AAC AGT CTT GGC GG-3’ and 5’-GGC GTA TCG CCA AAA TCA CC-3’, the Power SYBR Green PCR Master Mix (Applied Biosystems) and the relative standard curve quantitation method as implemented by the OneStep Plus Real-Time qPCR System (Applied Biosystems).

## Acknowledgements

Initial screening was carried out at Imperial College London Centre for Structural Biology EM facility. High resolution data were collected at the eBIC, Diamond Light Source (proposal EM14769). eBIC is funded by the Wellcome Trust, MRC and BBSRC. This work is funded by the BBSRC to XZ and MB (BB/N007816). XL is funded by a China Scholarship Council studentship. We are especially grateful to David Ratner for information concerning his experiments.

## Author contributions

XZ, DD and MB designed the studies. DD provided the initial Ocr samples. FY and XL prepared the samples. FY carried out competition experiments and performed the cryoEM anaysis. MJ conducted the in vitro transcription assays. IKL performed the in vivo experiments. XZ and MB wrote the manuscript with input from all the authors.

---

**Figure 1 – figure supplement 1.**
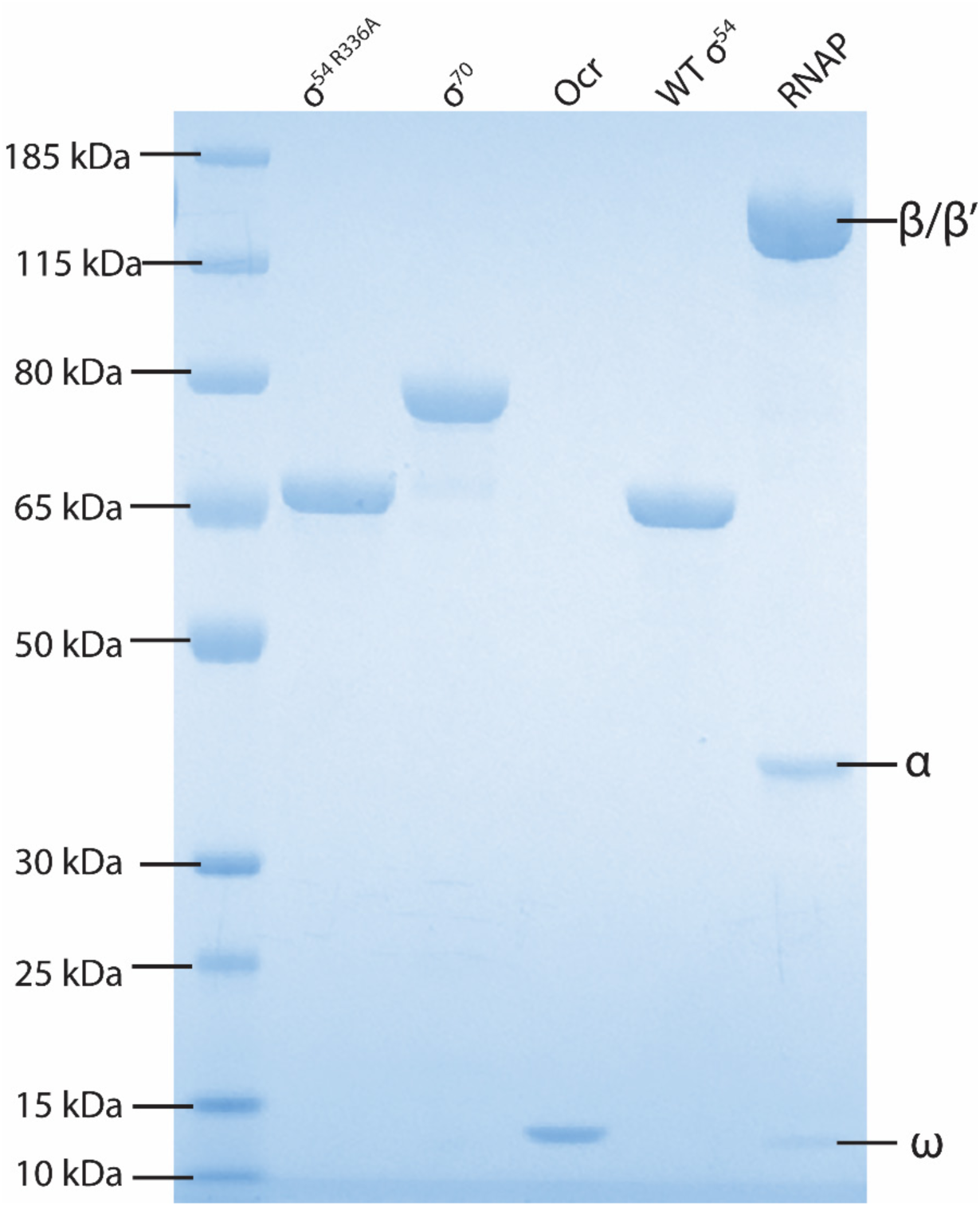
Quality of proteins used in this study as judged by SDS-PAGE gels.

**Figure 2 – figure supplement 1.**
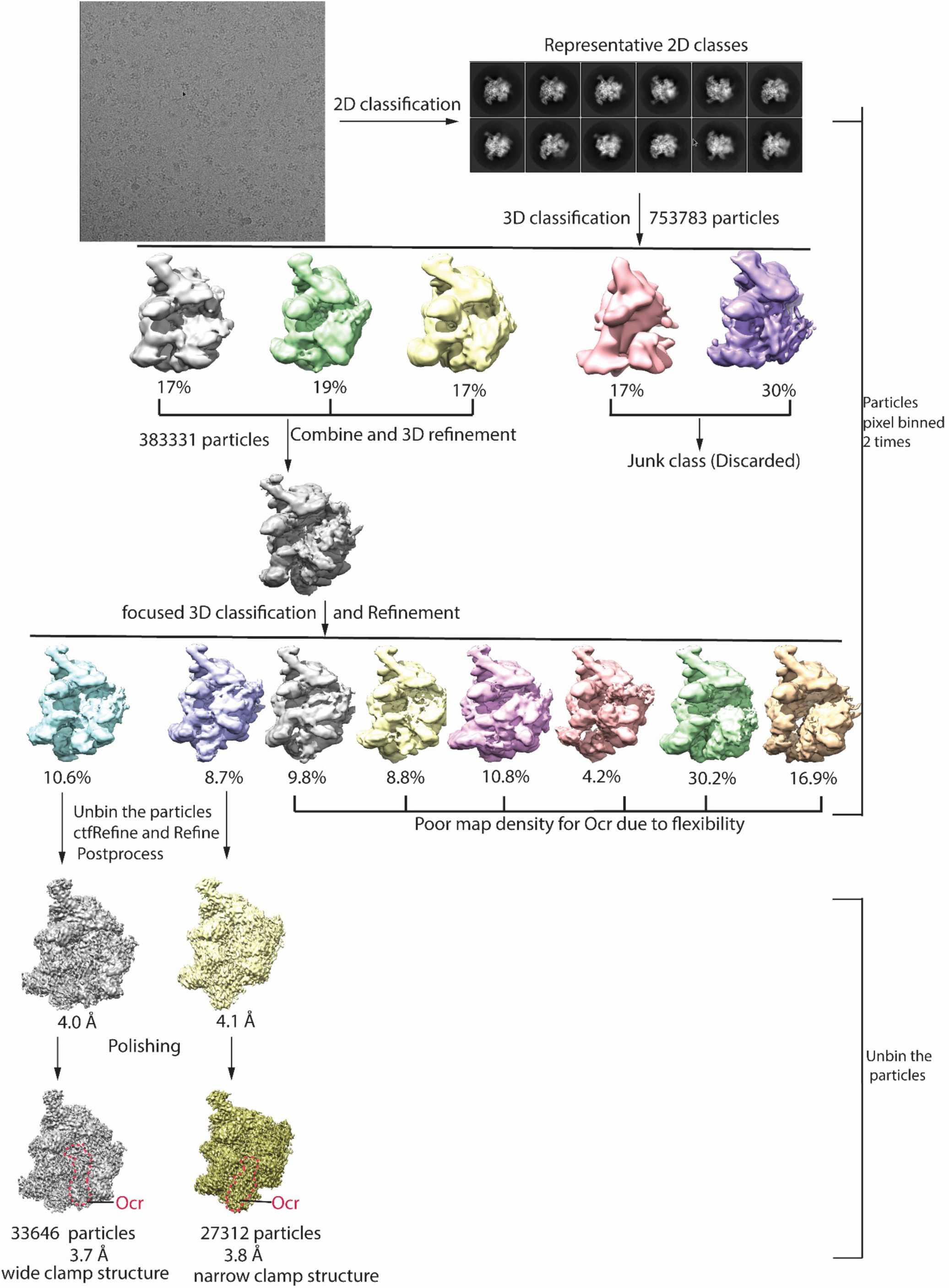
cryoEM data quality and Image processing flowchart. Included are a typical micrograph, representative 2D classes and image processing flowchart, and finally two distinct RNAP-Ocr complex obtained.

**Figure 2 – figure supplement 2.**
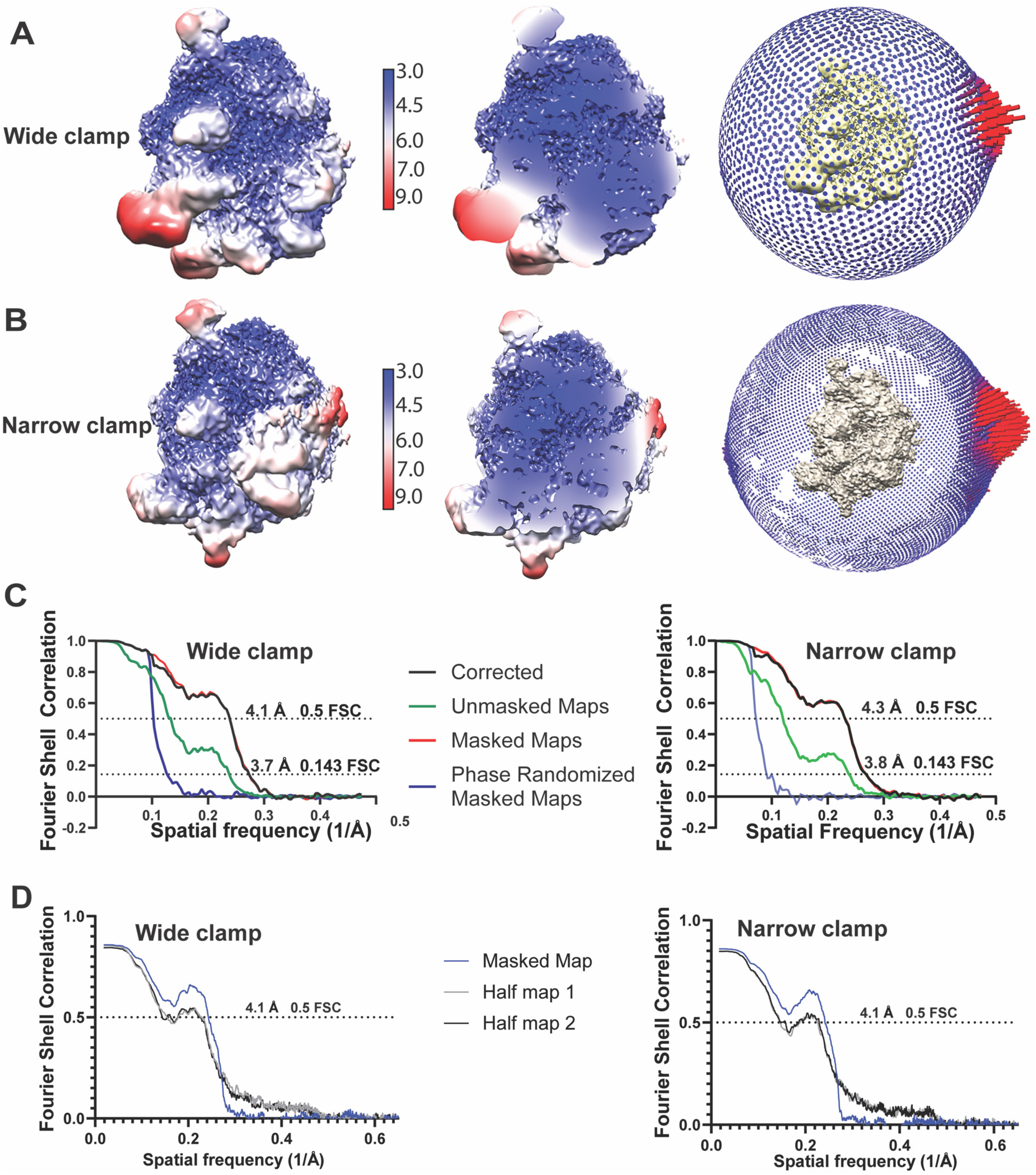
Quality of the two distinct RNAP-Ocr final 3D reconstructions. A) “wide-clamp” B) “narrow-clamp”. (A) and (B) shows local resolution maps in surface view and cut-through views (separated by resolution indicator), and angular distribution of particles (the extreme right panel of (A) and (B)). C) are the corresponding FSC curves of wide-clamp and narrow-clamp structures including corrected, unmasked, masked and phase randomized masked maps). D) FSC curves of model vs map for the two reconstructions.

**Figure 2 – figure supplement 3.**
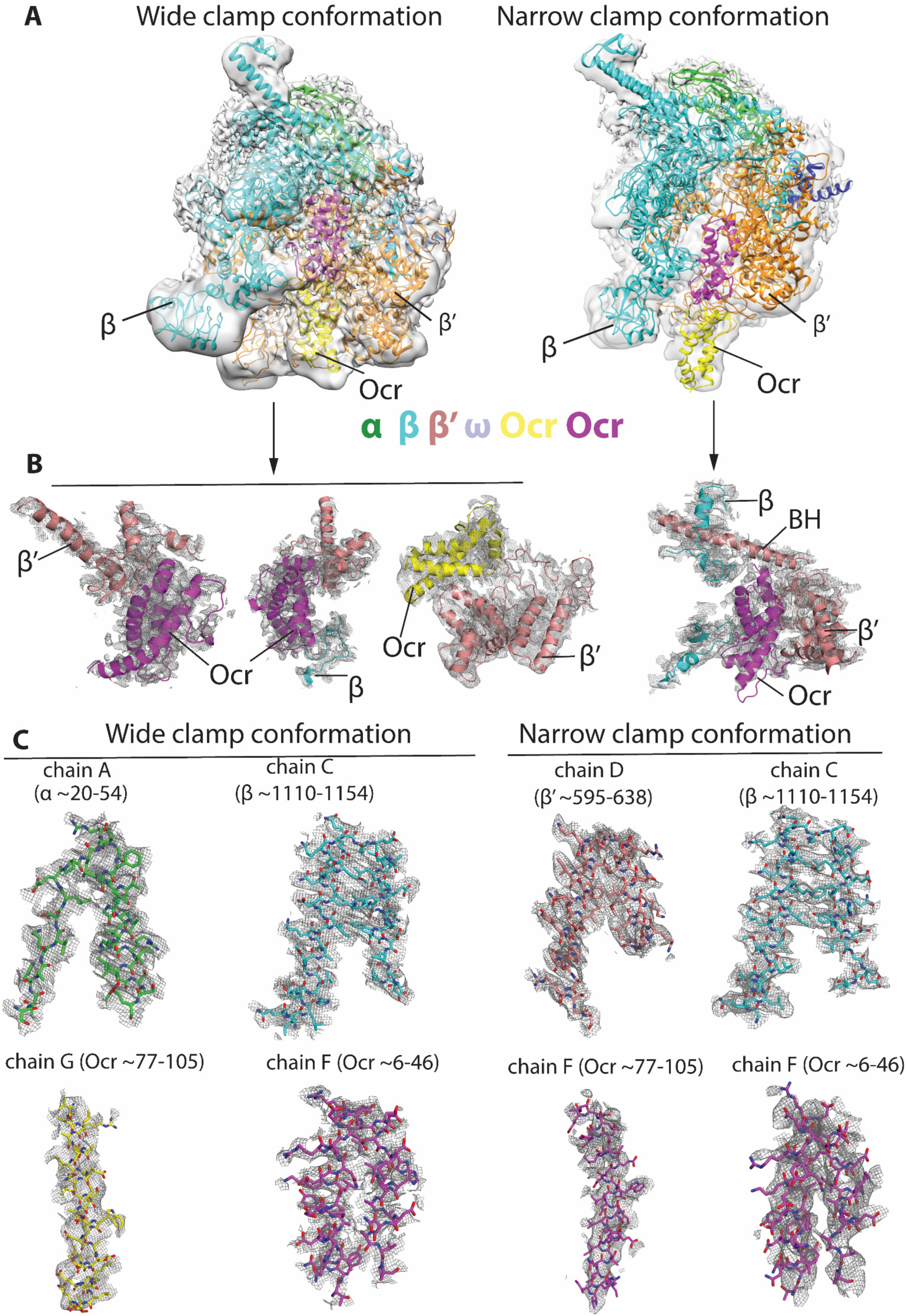
Representative electron density maps in the two reconstructions. **A)** showing overall fit of the atomic structural models into the corresponding reconstructions. Maps are filtered to local resolutions. left panel – wide clamp conformation (global resolution at 3.7 Å), right panels – narrow clamp conformation (global resolution at 3.8 Å). **B)** Views showing structural models of Ocr fitted into the electron density map and the surrounding RNAP. **C)**. Well defined regions showing clear density of RNAP and Ocr including side chains.

**Figure 5 – Table Supplement 1.**
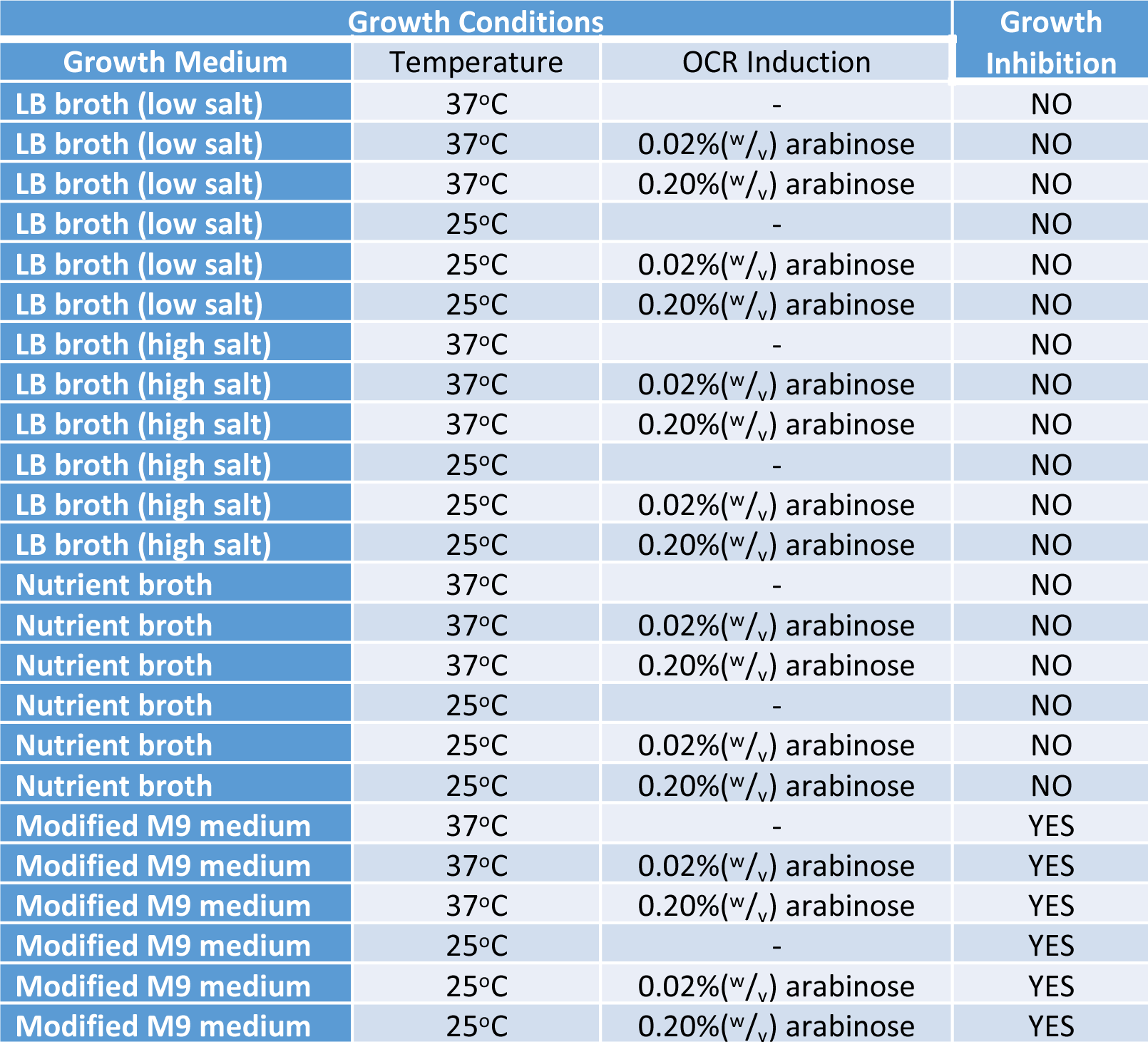
Pairwise comparisons of *E. coli* cells expressing Ocr (WT/pBAD18cm[*ocr*]) *versus* cells not expressing Ocr (WT/pBAD18cm).

**Figure 5 – Table Supplement 2.**
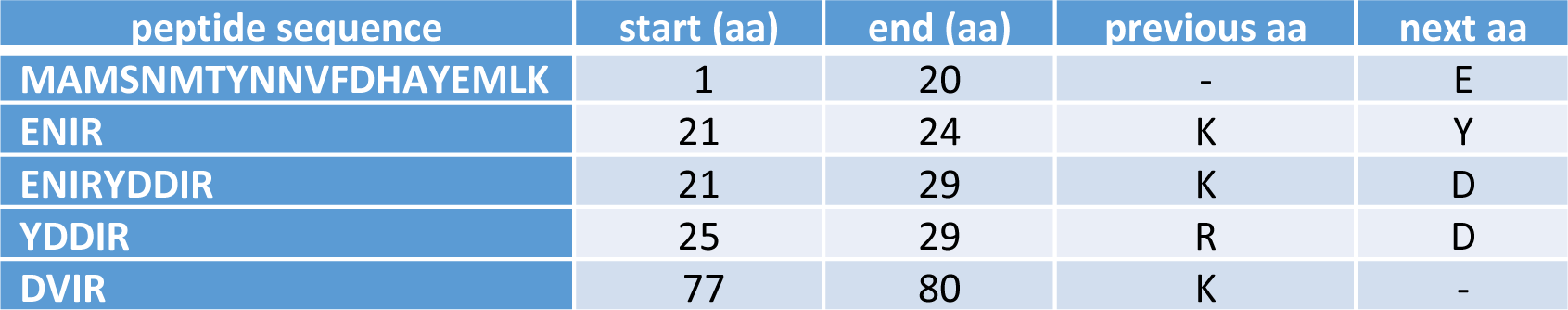
Tryptic peptides derived from Ocr following PMF of *E. coli* cell extracts.

**Figure 6 – figure supplement 1.**
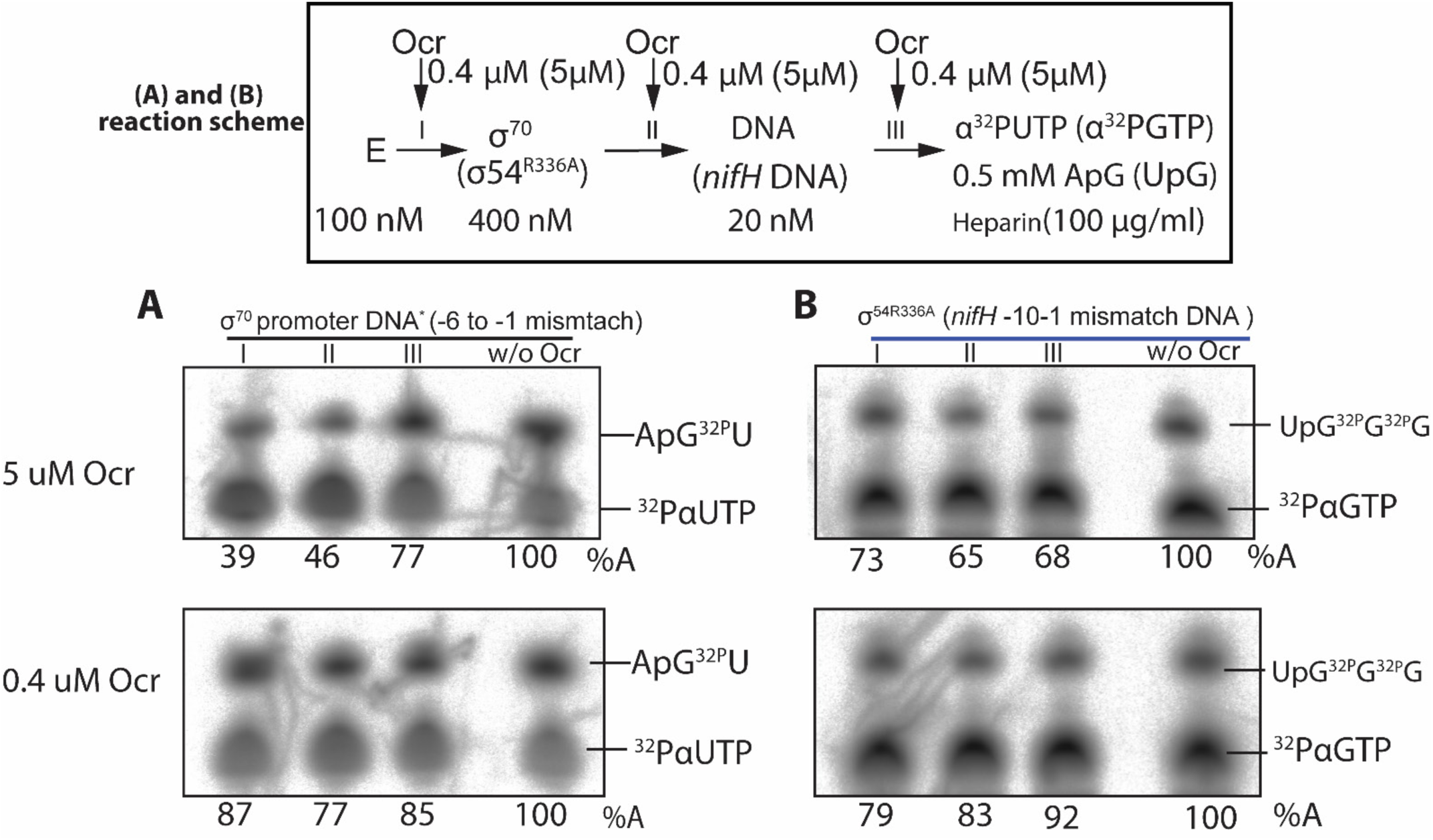
*In vitro* transcription assays. spRNA experiments on σ^70^ and σ^54^ using mismatch DNA (* Indicates that the DNA used here is the same as in Figure 7D – for open complex formation.)

**Figure 7 – figure supplement 1.**
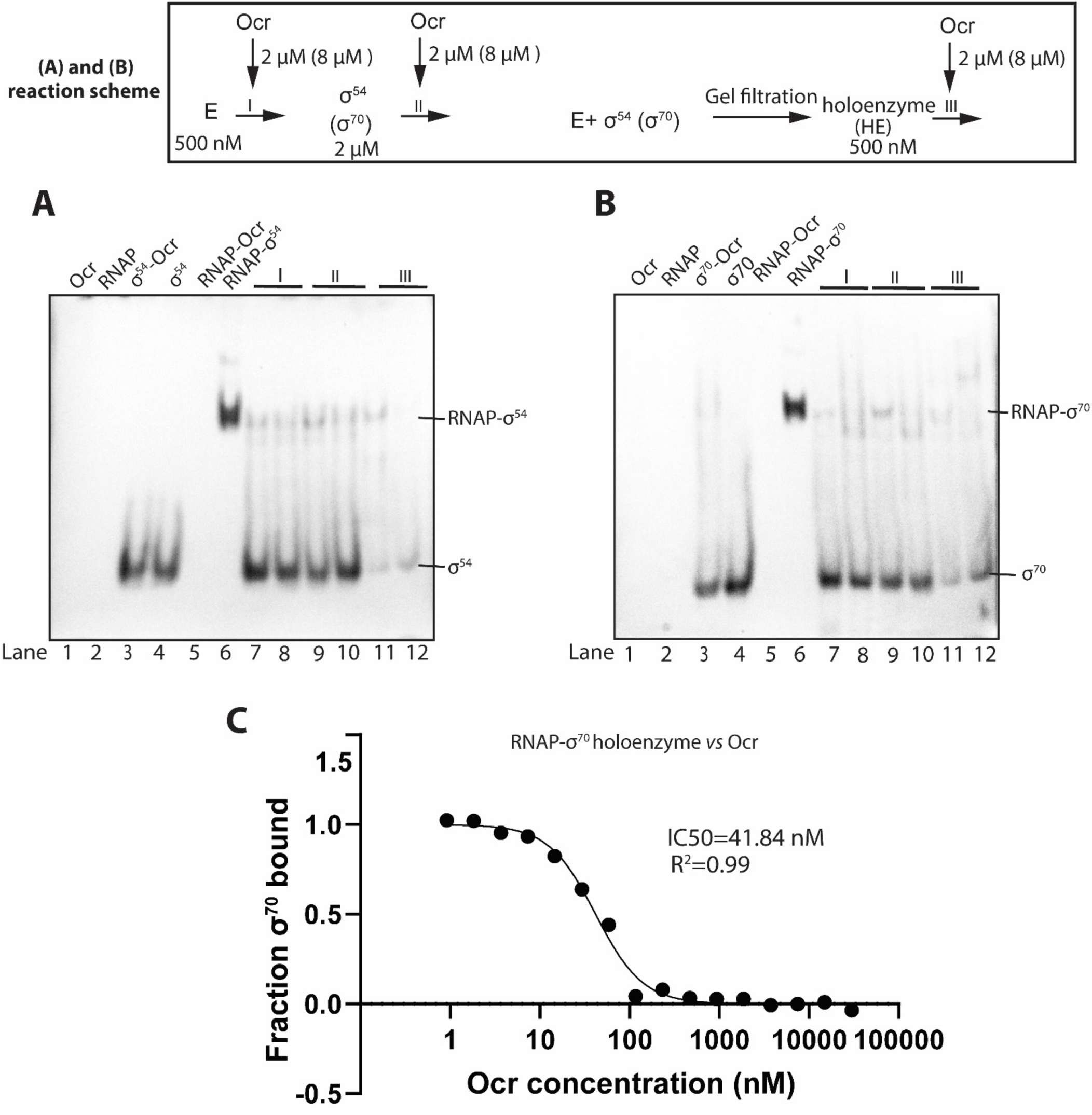
Ocr interferes with holoenzyme formation of σ^54^ and σ^70^, related to Fig. 5A-B. western blots against His-tags of σ^54^ and σ^70^ in the Native-PAGE gels as Fig.5A-B A) Ocr and σ^54^ holoenzyme formation. B) Ocr and σ^70^ holoenzyme formation. Reaction schemes are indicated above. E represents core enzyme RNAP, I, II, III indicate the point when Ocr is added during the reactions. Proteins and DNA concentrations are shown. In lanes 7, 9, 11. a 1:1 molar ratio of Ocr to σ was used, whereas in lane 8, 10, 12, a 4:1 molar ratio of Ocr to σ was used. C) Microscale thermophoresis (MST) experiment of Ocr binding to the RNAP-σ^70^ holoenzyme, supporting that Ocr competes with sigma in RNAP binding.

